# Boosting ATM Activity Promotes Longevity in Nematodes and Mice

**DOI:** 10.1101/240606

**Authors:** Minxian Qian, Zuojun Liu, Linyuan Peng, Fanbiao Meng, Xiaolong Tang, Ying Ao, Lei Shi, Mingyan Zhou, Ming Wang, Baoming Qin, Xinyue Cao, Zimei Wang, Zhongjun Zhou, Baohua Liu

**Author notes:** Minxian Qian and Zuojun Liu contributed equally to this work. Correspondence should be addressed to Dr Baohua Liu.

## Abstract

DNA damage accumulates with age^1^. However, whether and how robust DNA repair machinery promotes longevity is elusive. Here, we demonstrate that activation of ataxia-telangiectasia mutated (ATM) via low dose of chloroquine (CQ) promotes DNA damage clearance, rescues age-related metabolic shift, and extends lifespan in nematodes and mice. Molecularly, ATM phosphorylates SIRT6 deacetylase and thus prevents MDM2-mediated ubiquitination and proteasomal degradation. Extra copies of *Sirt6* in *Atm*-/- mice extend lifespan, accompanied with restored metabolic homeostasis. In a progeria mouse model with low ATM protein level and DNA repair capacity, the treatment with CQ ameliorates premature aging features and extends lifespan. Thus, our data highlights a pro-longevity role of ATM, for the first time establishing direct causal links between robust DNA repair machinery and longevity, and providing therapeutic strategy for progeria and age-related metabolic diseases.

## Introduction

A variety of metabolic insults frequently generate DNA lesions in mammalian cells, which, if wrongly repaired, may lead to somatic mutations and cell transformation^2^; if unrepaired, may accumulate and constantly activate DNA damage response (DDR), a unique feature and mechanism of senescence^3,4^. Ataxia telangiectasia mutated (ATM), a serine/threonine protein kinase, belongs to these key regulators of DDR^5^. Upon DNA damage, self-activated ATM phosphorylates downstream transducers and effectors, thus promoting DNA repair^6,7^. H2AX is one well-documented phosphorylation target of ATM; phosphorylated H2AX at S139 (γH2AX) is widely applied as a hallmark of DNA damage^8^. Accompanied with DNA repair function decline, γH2AX-enriched DNA damage foci accumulate in senescent cells and in tissues from aged animals^9^, supporting causal links between defective DDR and aging. In addition to γH2AX, CHK2, p53, SMC1, NBS1, BRCA1 and MDC1 are all ATM substrates^10^, of which phosphorylated Chk2 and p53 are key players of DDR-induced cell cycle arrest. In human fibroblasts, a dramatic decline of homologous recombination (HR) efficiency, attributable to defective recruitment of Rad51, was observed^11^. Similar defects in HR were also observed in Hutchinson-Gilford progeria syndrome (HGPS), which is predominantly caused by a *LMNA* G608G mutation^12^. In addition to DNA damage accumulation, inherited loss-of-function mutations in essential components of DNA repair machinery also accelerate aging in humans and mice^13^. Patients suffering from ataxia telangiectasia (A-T) develop prominent aging features in their second decades^14,15^. Werner syndrome, Bloom’s syndrome and Rothmund-Thomson syndrome are all progeria syndromes caused by mutations of genes that directly regulate DNA repair^16–19^. Homozygous disruption of *Atm* in mice recapitulates many premature aging features of A-T, like growth retardation, infertility, neurodegeneration, immunodeficiency and cancer predisposition^20^. Mouse models deficient in DNA repair elements, including DNA-PKcs, Ku70, Ku80, DNA ligase IV, Artemis or Ercc1, phenocopy premature aging features^21,22^. However, though tons of evidences support that defects in DNA repair accelerate ageing, whether and how robust DNA repair machinery promotes longevity is poorly understood.

Metabolic disturbance is another antagonistic hallmark of aging^23^. Although DNA repair deficiency is implicated in aging and age-related diseases including metabolic disorders^24,25^, the mechanistic linker between DNA repair machinery and metabolic reprogramming in aging is poorly understood. Notably, in response to oxidative stress, ATM phosphorylates Hsp27 thus to shift glucose metabolism from glycolysis to the pentose phosphate pathway (PPP)^26,27^. Inactivating ATM enhances glucose and glutamine consumption by inhibiting P53 and upregulating c-MYC^28^. The role of ATM in age-onset metabolic disturbances is yet unclear. On the other front, longevity-promoting genes, like NAD^+^-dependent sirtuins, are able to shunt energy metabolism away from anaerobic glycolysis toward TCA cycle^29,30^. Sirt6 cooperates with Hypoxia inducible factor-1α (HIF1α) via deacetylating transcription-active epigenetic marks to regulate glucose homeostasis^31^, and also modulates aerobic glycolysis^32^. Loss of *Sirt6* accelerates aging, accompanied with hypoglycemia and genomic instability^33,34^, whereas ectopic *Sirt6* inhibits insulin and insulin-like growth factor 1 (IGF1) signaling and extends lifespan in male mice^35^. Owing to up-regulated metabolic genes like glucose transporter GLUT-1, lactate dehydrogenase (LDH) and pyruvate dehydrogenase kinase (PDK), the glycolytic activity and lactate production were dramatically increased in *Sirt6*-/-cells and tumors^32^.

Here, we identified a progressive decline in ATM-centered DNA repair machinery during aging, along with shunted glucose metabolism to glycolysis. DNA damage-free activation of ATM by chloroquine (CQ) promotes DNA damage clearance, rescues the age-related metabolic shift, and extends lifespan in both nematodes and mice. Mechanistically, ATM phosphorylates and stabilizes pro-longevity protein SIRT6. Extra copies of *Sirt6* attenuate metabolic abnormality and extend lifespan in *Atm*-/- mice, and long-term treatment of CQ restores metabolic shift and extends lifespan in a progeria mouse model.

## Results

### ATM alleviates replicative senescence

In searching for genes/pathways that drive senescence, a gradual decline of ATM-centered DNA repair machinery was identified by RNAseq analysis (Supplementary Fig. 1a-e). Western blotting analysis confirmed progressively downregulated protein levels of ATM and its downstream target NBS1 and RAP80 in senescent human skin fibroblasts (HSFs) (Fig. 1a). Mouse embryonic fibroblasts (MEFs) with a limited growth capacity and senescent phenotypes when cultured *in vitro*^36–38^, and brain tissues from aged mice also showed progressive decline of ATM, NBS1, and PAP80 (Fig. 1b-c). Concomitantly, a reciprocal upregulation of γH2AX, indicating accumulated DNA damages, and p16^Ink4a^ level was observed in senescent HSFs, MEFs, and aged brain tissues (Fig. 1a-c). Knocking down *ATM* via shRNA accelerated senescence in HSFs, evidenced by increased β-galactosidase activity (Fig. 1d-e), enlarged morphology (Supplementary Fig.2a), accumulated γH2AX (Fig. 1f) and reduced cell proliferation (Supplementary Fig. 2b). Thus, ATM decline retards DDR and drives senescence.

**Figure 1:**
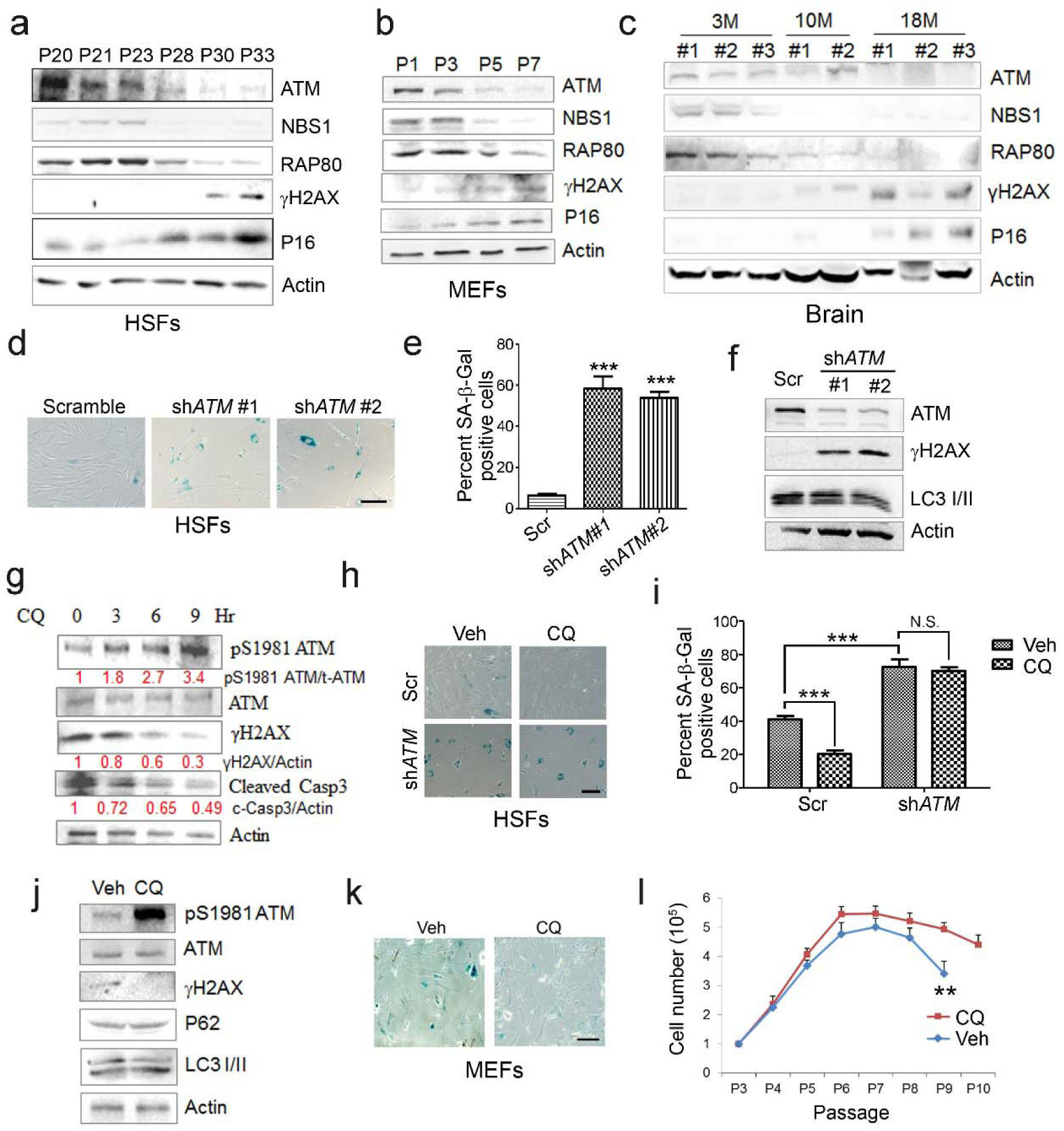
ATM activation by chloroquine alleviated senescence. (**a**) Immunoblots showing protein levels of ATM, NBS1 and RAP80 in human skin fibroblasts (HSFs). Gradually increased level of p16 indicates cellular senescence. Elevated γH2AX level indicates accumulated DNA damage. (**b**) Immunoblots showing protein levels of ATM, NBS1 and RAP80 in mouse embryonic fibroblasts (MEFs). (**c**) Immunoblots showing protein levels of ATM, NBS1 and RAP80 in brain tissues isolated from 3, 10 and 18-month-old male mice. (**d**) SA-β-Gal staining in HSFs treated with sh-*ATM* or scramble shRNA. Scale bar, 100 µm. (**e**) Quantification of SA-β-Gal-positive staining of (**d**) from five views randomly captured for each group. Data represent mean ± SEM. *** *P* < 0.001. (**f**) Immunoblots showing increased γH2AX and unaffected LC3I/II in HSFs treated with sh-*ATM* or scramble shRNA. (**g**) Immunoblots showing protein levels of pS1981 ATM, γH2AX and cleaved caspase-3 in HSFs treated with 10 μM of CQ for indicated time. (**h**) SA-β-Gal staining in HSFs expressing either scramble or *ATM* shRNA treated with 1 μM CQ or DMSO (12 h). Scale bar, 100 µm. (**i**) Quantification of SA-β-Gal-positive staining of (h) from five views randomly captured for each group. Data represent mean ± SEM. \*\*\**P* < 0.001; ‘N.S.’ indicates no significant difference. (**j**) Immunoblots showing protein levels of γH2AX, p62 and LC3 in MEFs treated with 1 μM CQ or DMSO. Noted that the levels of p62 and LC3 were merely affected. (**k**) SA-β-Gal staining in primary MEFs treated with 1 μM CQ or DMSO. Scale bar, 100 µm. (**l**) Cumulative cell numbers of continuously cultured primary MEFs treated with 1 μM CQ or DMSO. Data represent mean ± SEM. \*\**P* < 0.01.

Other than DNA damage, ATM can be activated by chloroquine (CQ), an antimalarial drug that modulates chromatin confirmation^6^. Indeed, we found that low dose of CQ increased pS1981 auto-phosphorylation of ATM but not γH2AX (Supplementary Fig. 2c). We then investigated whether activating ATM by CQ ameliorates senescence. As shown, the CQ treatment activated ATM (pS1981), promoted clearance of DNA damage (γH2AX), and inhibited apoptosis (cleaved Casp3) in HSFs (Fig. 1g). Also, the CQ treatment suppressed β-galactosidase activity, but that was abrogated when *ATM* was knocked down (Fig. 1h-i). Likewise, the CQ treatment activated Atm, cleared up accumulated DNA damage (Fig. 1j), suppressed β-galactosidase activity (Fig. 1k and Supplementary Fig. 2d), and prolonged replicative lifespan in MEFs (Fig. 1l). Despite that both doses of 10 μM and 1 μM CQ activated ATM, the dose-dependent toxicity assay showed CQ in 1 μM is suitable for long-term treatment (Supplementary Fig. 2e-f). Of note, *ATM* KD or low dose of CQ applied in this study had little effect on basal autophagic activity (Fig. 1f, j and Supplementary Fig. 2g). Collectively, CQ activates ATM to alleviate replicative senescence.

### An ATM-SIRT6 axis underlies age-associated metabolic reprogramming

A-T patients lacking functional ATM display features of premature aging, accompanied with insulin resistance and glucose intolerance^39,40^. Senescent cells exhibit enhanced glycolysis but impaired mitochondrial respiration, and increased lactate for ATP generation^41,42^. As such, we must ask whether ATM decline triggers age-associated metabolic shift. Indeed, glycolytic gene *LDHB* and *PDK1* were dramatically increased in senescent MEFs and HSFs (Fig. 2a and Supplementary Fig. 3a), and in liver tissues from *Atm*-/- mice (Supplementary Fig. 3b). Significantly, activating ATM via CQ suppressed senescence-associated glycolysis (Fig. 2a and Supplementary Fig. 3a). The inhibitory effect on glycolysis was diminished when *ATM* was depleted in HepG2 cells (Fig. 2b), suggesting the role of ATM in inhibiting glycolysis.

**Figure 2:**
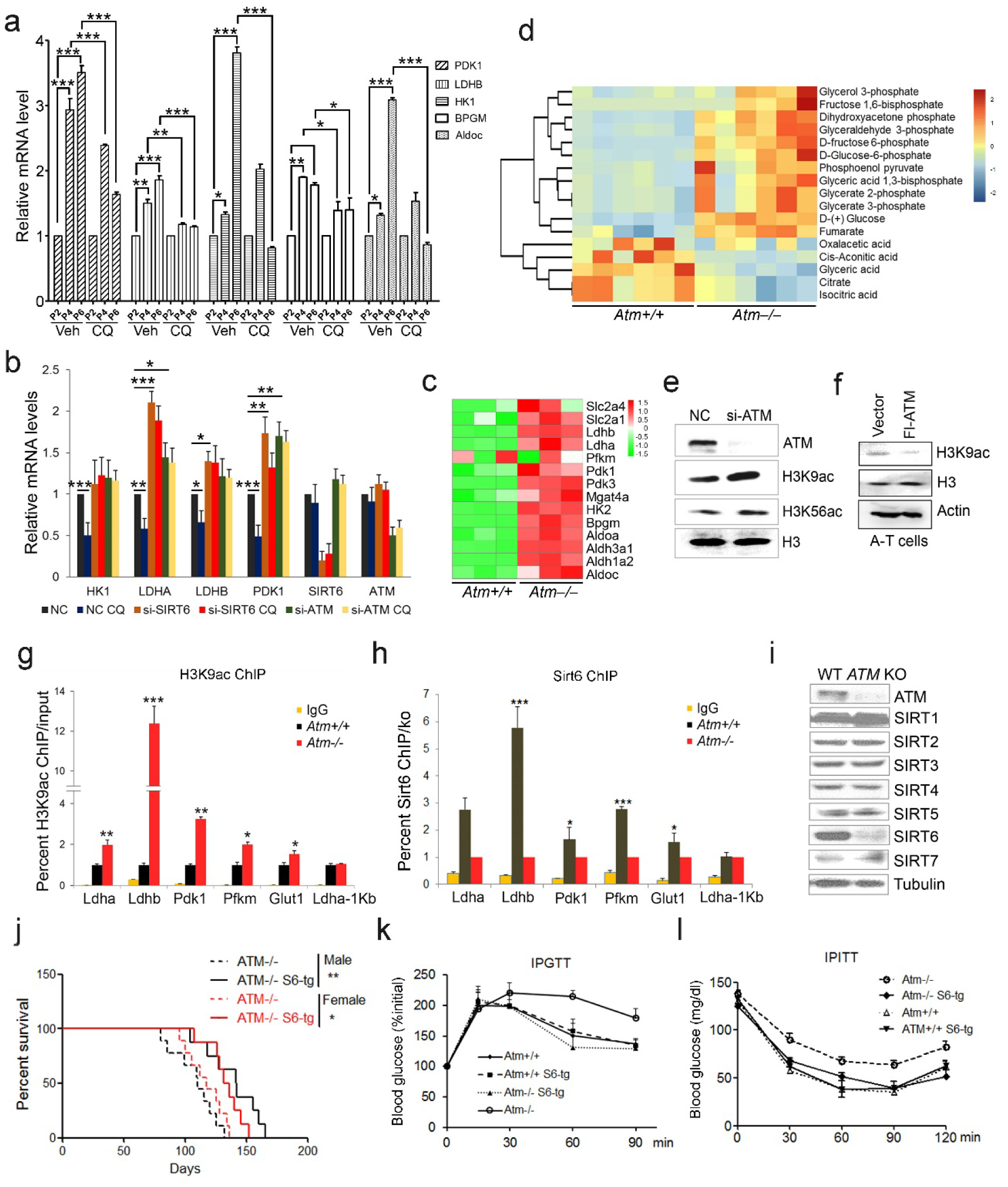
ATM-SIRT6 axis regulates age-related metabolic reprogramming. (**a**) Quantitative RT-PCR analysis of mRNA levels of indicated glycolytic genes in different passages of MEFs with or without treatment of CQ. Data represent mean ± SEM. \**P* < 0.05, \*\**P* < 0.01, \*\*\**P* < 0.001. (**b**) Quantitative RT-PCR analysis of mRNA levels of indicated glycolytic genes in Scramble (NC), si-*SIRT6* or si-*ATM* HepG2 cells incubated with or without CQ (10μM, 6 h). Data represent mean ± SEM. \**P* < 0.05, \*\**P* < 0.01, \*\*\**P* < 0.001. (**c**) Heatmap representation of RNA-Seq data showing relative changes of glycolytic genes in *Atm*-/- MEF cells. Red and green indicate up- and downregulation, respectively. (**d**) Heatmap showing relative levels of metabolites in *Atm*+/+ and *Atm*-/- MEF cells of p53 null background, analyzed by LC-MS. Red and blue indicate up-and downregulation, respectively. (**e**) Immunoblots showing protein levels of H3K9ac and H3K56ac in *ATM*-deficient HepG2 cells. (**f**) Immunoblots showing levels of H3K9ac in A-T cells reconstituted with Flag-ATM. (**g**-**h**) ChIP analysis showing enrichment of H3K9ac and SIRT6 at the promoter regions of indicated genes in *Atm*+/+ and *Atm*-/- MEFs. Data represent mean ± SEM of 3 independent experiments. \**P* < 0.05, \*\**P* < 0.01, \*\*\**P* < 0.001. (**i**) Immunoblots showing protein levels of sirtuins in wild-type (WT) and *ATM* knockout (KO) HEK293 cells. (**j**) Kaplan-Meier survival of *Atm*-/- and *Atm-/-; Sirt6*-Tg male (n = 11 in each group) and female (n = 9 in each group) mice. \*\**P* < 0.01. (**k**) Results of glucose tolerance tests in *Atm*+/+, *Atm*-/-, and *Atm*-/-; *Sirt6*-Tg mice. Data represent mean ± SEM, n = 6. \*\**P* < 0.01, \*\*\**P* < 0.001. (**l**) Results of insulin tolerance tests in *Atm*+/+, *Atm*-/-, and *Atm-/-; Sirt6*-Tg mice. Data represent mean ±SEM, n = 6. \*\**P* < 0.01.

To address how ATM regulates glycolysis, we performed RNA-Seq in *Atm*-/- MEF cells, and revealed a significant upregulation of glycolytic pathways (Fig. 2c). Specific genes were validated by q-PCR (Supplementary Fig. 3c). As p53 is critical in glycolysis^43,44^, we further analyzed metabolomics of *Atm*-/- and control MEF cells in *p53* null background. As shown, the metabolic profile exhibited a clear shift, i.e. mitochondrial electron transport chain and intermediates of TCA cycle were reduced, while intermediates of glycolysis were elevated (Fig. 2d, Supplementary Fig. 3d-e and Table S1). The data suggest *ATM* deficiency enhances anaerobic glycolysis in a p53-independent manner.

Sirt6 deacylase is able to shunt energy metabolism away from anaerobic glycolysis to TCA cycle via H3K9ac-mediated local chromatin remodeling^31,32^. We noted the level of H3K9ac was enhanced in cells depleted *ATM* (Fig. 2e). Re-expressed ATM in A-T cells suppressed H3K9ac level (Fig. 2f). Further ChIP analysis showed that H3K9ac was enriched at the promoter regions of glycolytic genes in *Atm*-/- cells (Fig. 2g), where the relative occupancy of SIRT6 was abolished (Fig. 2h). Consistent with increased H3K9ac level, SIRT6 protein level was dramatically downregulated in *Atm*-/- mouse livers, and *ATM* deficient HepG2, U2OS, and HEK293 cells (Supplementary Fig. 3f-i). In contrast, other sirtuins were merely affected (Fig. 2i), and mRNA levels of all sirtuins remained unchanged (Supplementary Fig. 3j). Moreover, transcriptomic analysis and q-PCR data illustrated that *Sirt6* depletion upregulated a similar cluster of genes essential for glycolysis (Supplementary Fig. 4a-b and Table S2). More importantly, the hyper-activated glycolytic pathway caused by *ATM* deficiency was completely restored by ectopic *SIRT6* in HepG2 cells (Supplementary Fig. 4c). The CQ treatment upregulated SIRT6 level and reduced H3K9ac level, especially at regulatory regions of glycolytic genes (Supplementary Fig. 4d-e). Knocking down *SIRT6* abolished the inhibitory effect of CQ on glycolysis (Fig. 2b). Though levels of SIRT1 and SIRT7 were also declined along with senescence, they were not restored by CQ treatment (Supplementary Fig. 4f). Additionally, *ATM* depletion in HEK293 cells, HSFs and MEFs, significantly downregulated SIRT6 protein level, with no effect on SIRT1 or SIRT7 protein level (Fig. 2i and Supplementary Fig. 4g-h). Thus, these data suggest ATM decline triggers age-associated metabolic shift via SIRT6-mediated chromatin remodeling.

Other than metabolic abnormality, depleting *Sirt6* leads to various premature aging features and shortened lifespan^45^, whereas extra copies of *Sirt6* promotes longevity in male mice^35^. Given that Sirt6 was destabilized in *Atm* null mice, we asked whether *Sirt6* transgene could rescue premature aging phenotypes and shortened lifespan in *Atm*-/- mice. To this end, we generated *Sirt6* transgenic mice by microinjection, and bred them with *Atm*-/- mice. The overexpression of Sirt6 was demonstrated by Western blotting (Supplementary Fig. 4i). Significantly, ectopic *Sirt6* restored the elevation of serum lactate and blood glucose levels, and extended lifespan of *Atm*-/- mice in both genders (Fig. 2j and Supplementary Fig. 4j-k). Importantly, *Atm-/-; Sirt6-*tg mice exhibited improved glucose tolerance and decreased insulin resistance (Fig. 2k-l). Given little difference was observed in glucose metabolism between young WT and Sirt6-transgenic mice^35^, these data suggest a contributing role of the Atm-Sirt6 axis in the age-associated metabolic reprogramming.

### ATM phosphorylates and stabilizes SIRT6

Next, we examined how ATM regulates SIRT6. Significantly, the overexpression of *ATM* increased SIRT6 level, but that was abolished when ATM was S1981A-mutated (Fig. 3a). Of note, the S1981A mutation blocks dimeric ATM dissociation and therefore the ATM activation^6^. Moreover, in addition to CQ, hypotonic buffer (20 mM NaCl), low glucose (LG), DNA-damaging agent CPT and doxorubicin (Dox) all activated ATM and concomitantly increased SIRT6 protein level (Supplementary Fig. 5a-c), but this effect was abrogated in *ATM*-depleted cells (Supplementary Fig. 5b-c). These data implicate a direct regulation of SIRT6 stability by ATM kinase activity. To confirm it, we first performed co-immunoprecipitation (Co-IP) in cells transfected with various FLAG-sirtuins. Very interestingly, ATM was predominantly associated with SIRT6 among seven sirtuins (Fig. 3b). The interaction was further confirmed at both ectopic and endogenous levels (Fig. 3c and Supplementary Fig. 5d). Immunofluorescence microscopy showed co-localization of SIRT6 and ATM protein in the nucleus (Fig. 3d). Domain mapping experiment indicated that the C-terminal domain was required for SIRT6 binding to ATM (Supplementary Fig. 5e). To determine whether ATM physically binds to SIRT6, 10 consecutive recombinant GST-ATM proteins were obtained and the binding to purified His-SIRT6 was analyzed. As shown, His-SIRT6 strongly bond to GST-ATM-4 (residues 770–1102) (Fig. 3e), the N-terminal HEAT repeat domain of ATM, which are predicted to bind to phosphorylation substrates, such as NBS1^46^.

**Figure 3:**
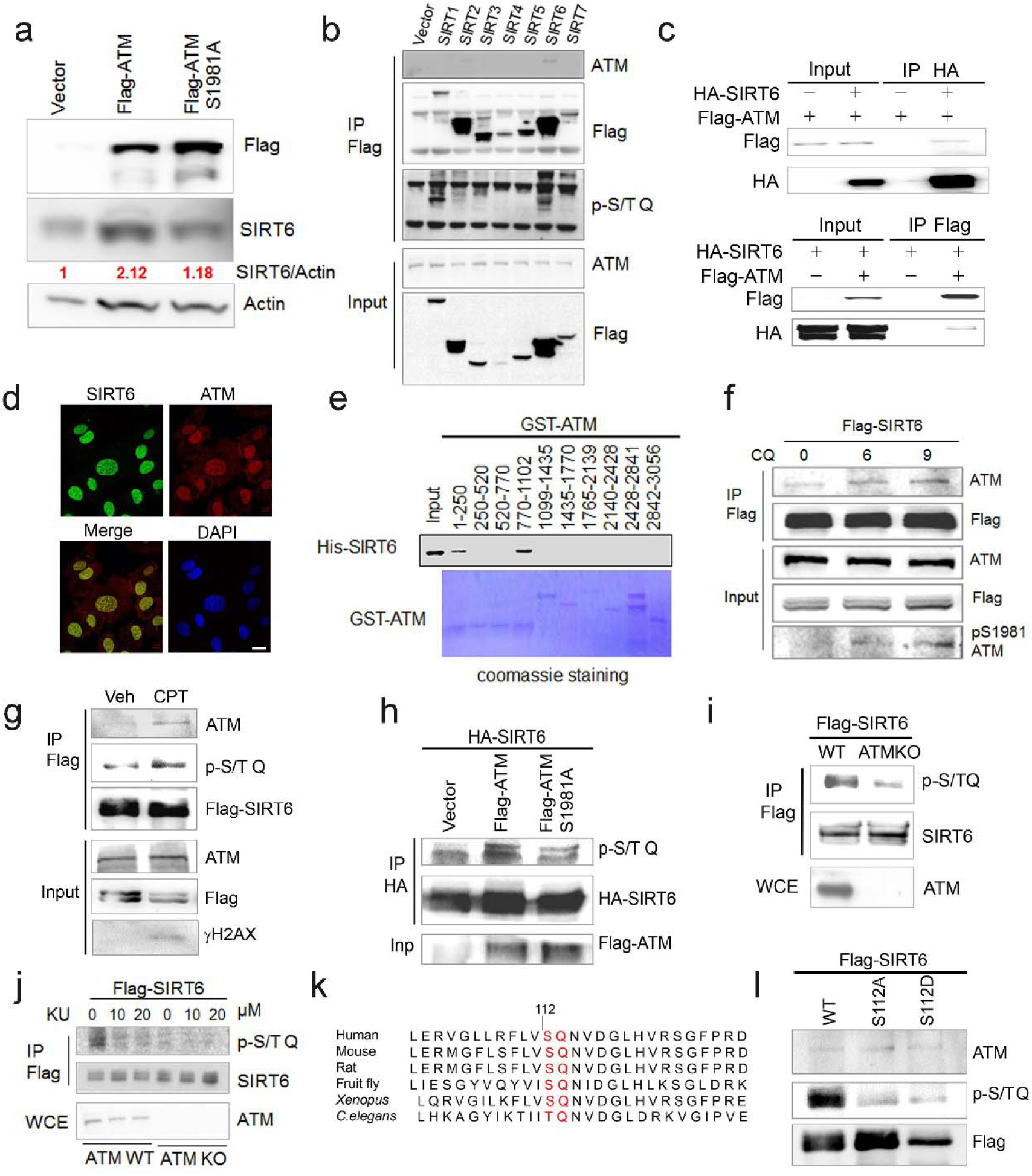
ATM interacts with and phosphorylates SIRT6. (**a**) Immunoblots showing protein levels of SIRT6 in HEK293 cells expressing Flag-ATM or Flag-ATM S1981A. (**b**) Immunoblots showing endogenous ATM and p-S/TQ motif in anti-Flag immunoprecipitates in HEK293 cells transfected with empty vector or Flag-sirtuins. (**c**) Immunoblots showing Flag-ATM and HA-SIRT6 in anti-HA (upper) or anti-Flag (lower) immunoprecipitates in HEK293 cells transfected with indicated constructs. (**d**) Representative photos of immunofluorescence staining of SIRT6 and ATM in U2OS cells, showing the co-localization in nucleus. Scale bar, 50 µm. (**e**) GST pull-down assay showing bacterially expressed His-SIRT6 predominantly bound to GST-ATM fragment 4 (770–1102), the N-terminal HEAT-repeat of ATM. (**f**) Immunoblots showing the increased binding capacity of ATM and SIRT6 under the treatment of (10 µM) CQ for the indicated time. (**g**) Immunoblots showing ATM and p-S/TQ in anti-Flag immunoprecipitates in HEK293 cells expressing Flag-SIRT6 treated with CPT (0.4 µM) or DMSO. (**h**) Immunoblots showing level of p-S/TQ SIRT6 in HEK293 cells co-transfected with HA-SIRT6 and Flag-ATM, Flag-ATM S1981A, or empty vector. (**i**) Immunoblots showing p-S/T Q SIRT6 in WT or ATM KO HEK293 cells. (**j**) Immunoblots showing p-S/T Q level of SIRT6 in *ATM* WT or KO HEK293 cells treated with DMSO and KU55933 (10 or 20 µM, 2 h). (**k**) Alignment of protein sequence of human SIRT6 and orthologues in mouse, rat, fruit fly, *Xenopus*, and *C. elegans*. A conserved S^112^ Q^113^ motif was highlighted. (**l**) Immunoblots showing p-S/T Q level of Flag-SIRT6, Flag-SIRT6 S112A, or Flag-SIRT6 S112D in HEK293 cells.

We next examined whether ATM phosphorylates SIRT6. Firstly, we found the CQ or CPT treatment significantly enhanced the binding of SIRT6 to ATM (Fig. 3f and Supplementary Fig. 5f), whereas the S1981A mutant blocked such association (Supplementary Fig. 5g). ATM preferentially phosphorylates S/T-Q motif. In the presence of CPT, the increased p-S/TQ level of SIRT6 was identified (Fig. 3g). Of note, lambda protein phosphatase (λPP) diminished p-S/TQ level of SIRT6 (Supplementary Fig. 5h). Likewise, the p-S/TQ level of SIRT6 was elevated in cells treated with low glucose, which activates ATM by ROS generation^47,48^ (Supplementary Fig. 5i). Moreover, ectopic ATM significantly increased p-S/TQ level of SIRT6, and that was abolished in case of S1981A-mutated (Fig. 3h). Consistently, a pronounced reduction of p-S/TQ level of SIRT6 was observed in cells lacking *ATM* or treated with KU55933, a selective and specific ATM kinase inhibitor^49,50^ (Fig. 3i-j). The decrease in p-S/TQ level was primarily attributable to loss of *ATM*, as it was restored by ectopic FLAG-ATM in a dose-dependent manner (Supplementary Fig. 5j). Indeed, SIRT6 has one evolutionarily conserved S^112^Q^113^ motif (Fig. 3k). We therefore constructed S112A and S112D mutations, which resembles hypo- and hyper-phosphorylated form respectively. As shown, these mutations almost abolished the pS/T-Q level of FLAG-SIRT6 (Fig. 3l). Further *in vitro* kinase assay showed that ATM could phosphorylate GST-SIRT6, but not S112A mutant (Supplementary Fig. 5k). Collectively, the data suggest that ATM directly phosphorylates SIRT6 at Serine 112.

We next examined whether ATM is involved in regulating SIRT6 protein stability. Notably, compared to wild-type or vehicle control, the degradation rates of ectopic and endogenous SIRT6 were largely increased in *ATM* KO HEK293 cells, *Atm***-/-** MEFs, or cells incubated with ATM specific inhibitor KU55933 in the presence of cycloheximide (CHX) (Fig. 4a-b and Supplementary Fig. 6a-c). Recently MDM2 was demonstrated to ubiquitinate SIRT6 and promote its proteasomal degradation^51^. We therefore examined the polyubiquitination level of SIRT6. As shown, the ubiquitination level of FLAG-SIRT6 in *ATM* KO cells was significantly elevated compared with wild-type (Supplementary Fig. 6d). While the S112A mutant markedly enhanced poly-ubiquitination level of SIRT6, the S112D merely affected it (Supplementary Fig. 6e). Moreover, the S112A accelerated SIRT6 degradation, whereas the S112D retarded it (Fig. 4c-d), indicating that the Ser112 phosphorylation by ATM regulates SIRT6 ubiquitination and thus protein stability. Indeed, ectopic MDM2 enhanced polyubiquitination level of FLAG-SIRT6 (Supplementary Fig. 6f). In case of *ATM* depleted or SIRT6 S112A-mutated, the binding capacity of SIRT6 to MDM2 was enhanced (Fig. 4e and Supplementary Fig. 6g). In searching for key residues that are poly-ubiquitinated by MDM2, we identified two clusters of lysine residues, i.e. K143/145 and K346/349, which are conserved across species. We then generated KR mutations of these residues, and found K346/349R remarkably reduced the poly-ubiquitination level of SIRT6 (Supplementary Fig. 6h). Individual KR point mutation showed that K346R significantly blocked MDM2-mediated ubiquitination and degradation of SIRT6, whereas K349R hardly affected it (Fig. 4f-g). More importantly, the K346R restored the increased ubiquitination and accelerated protein degradation of SIRT6 S112A (Supplementary Fig. 6i-j). Collectively, these data indicate that K346 is subjected to MDM2-mediated ubiquitination and is responsible for the degradation of SIRT6, which is inhibited by ATM-mediated S112 phosphorylation.

**Figure 4:**
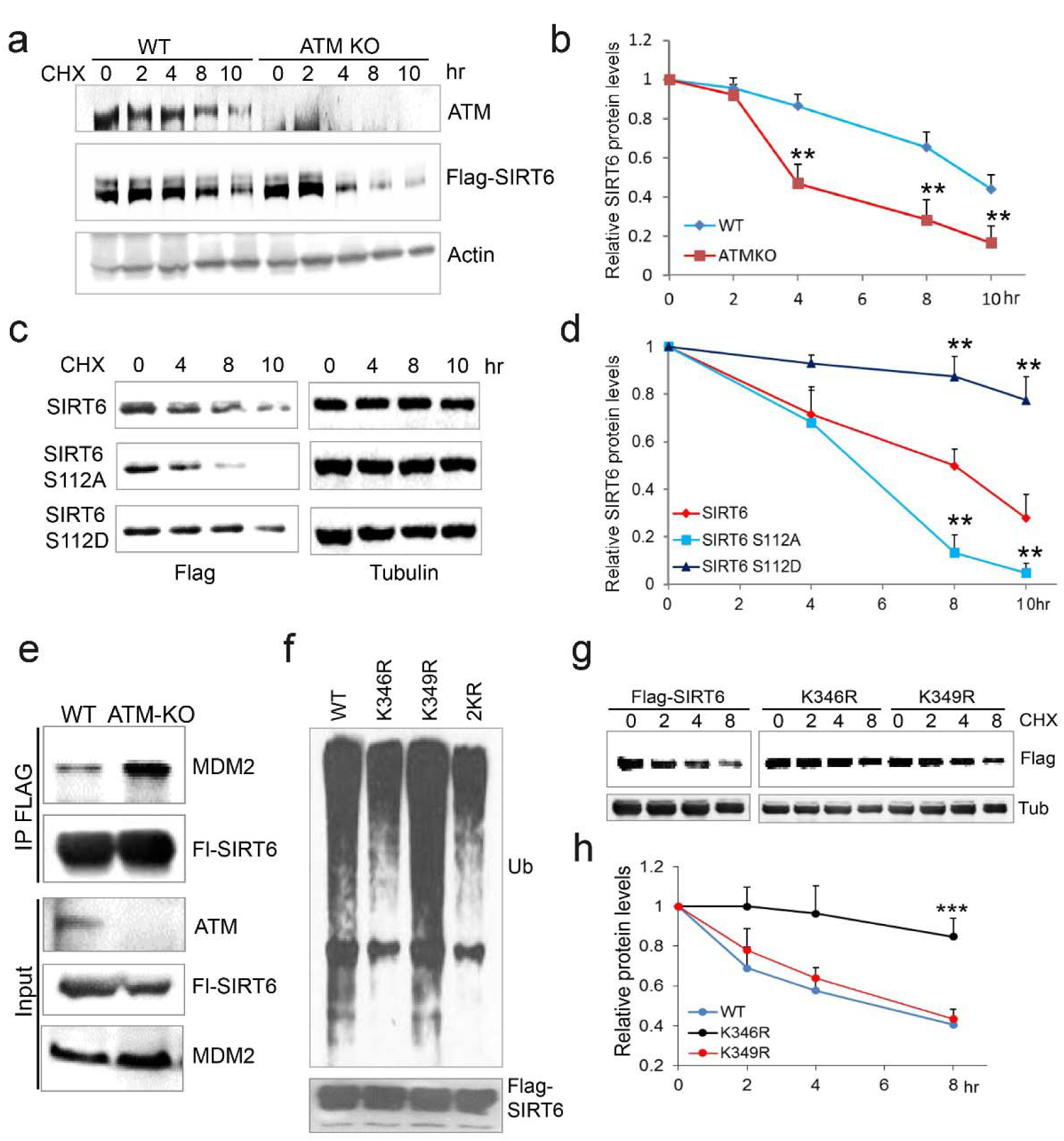
ATM prevents ubiquitination and degradation of SIRT6. (**a**) Immunoblots showing protein levels of Flag-SIRT6 in WT and *ATM* KO HEK293 cells treated with CHX (50 µg/ml) for indicated periods of time. (**b**) Quantification of protein levels in (**a**) by ImageJ^®^. Data represent mean ± SEM of three independent experiments. \*\**P* < 0.01. (**c**) Immunoblots showing protein levels of Flag-SIRT6, S112A and S112D in the presence of CHX (50 μg/ml) for indicated periods of time. (**d**) Quantification of protein levels in (**c**) by ImageJ^®^. Data represent mean ± SEM of three independent experiments. \*\*\**P* < 0.001. (**e**) Immunoblots showing increased binding capacity between SIRT6 and MDM2 in *ATM* KO HEK293 cells. (**f**) Immunoblots showing ubiquitination of Flag-SIRT6, K346R, K349R and K346/349R (2KR) in HEK293 cells. Noted that 2KR and K346R abrogated the ubiquitination of Flag-SIRT6. (**g**-**h**) Upper, immunoblots showing protein levels of Flag-SIRT6, K346R and K349R in the presence of CHX (50 μg/ml) for indicated periods of time. Lower, quantification of protein levels by ImageJ^®^. Data represent mean ± SEM of three independent experiments. \*\*\**P* < 0.001.

### Activating ATM via CQ promotes longevity

The cellular data suggest a pro-longevity function of ATM. We then tested it at organismal level. We employed *Caenorhabditis elegans*, which have a short lifespan of approximate 30 days. Nematodes deficient for *atm-1*, an orthologue of mammalian *ATM*, and wild-types were exposed to various doses of CQ (see Materials and Methods). Significantly, the period treatment with CQ (1.0 µM) extended the median lifespan (~14%) of *C. elegans* (Fig. 5a). The lifespan-extending effect was abolished in *atm-1* KO (Fig. 5b) or in SIRT6 homolog *sir-2.4* KD nematodes (Supplementary Fig. 7a-b). The data suggest that CQ promotes longevity in an ATM- and SIRT6-dependent manner. We further examined the beneficial effect of CQ in a HGPS model, i.e. *Zmpste24***-/-** mice, which has a shortened lifespan of 4–6 months^52^ and impaired ATM-mediated DNA repair signaling^53^. Here we found, Atm level was dramatically reduced in *Zmpste24***-/-** cells and tissues (Supplementary Fig. 7c-d). Significantly, the CQ treatment activated Atm, stabilized Sirt6, decreased the accumulated DNA damage, inhibited glycolysis, and alleviated senescence in *Zmpste24***-/-** cells (Fig. 5c-d and Supplementary Fig. 7e-f). The CQ treatment also held off body weight decline, increased running endurance, and prolonged lifespan in *Zmpste24***-/-** mice (Fig. 5e-f and Supplementary Fig. 7g). We then extended the study to physiologically aged mice. In this case, 12-month-old “old” male mice were intraperitoneally administrated with lose dose of CQ (3.5mg/kg) twice a week. Remarkably, compared to the saline-treated control group, the CQ treatment inhibited glycolysis, lowered down serum lactate level, attenuated body weight decline (Fig. 5g-h and Supplementary Fig. 7h). Most remarkably, the CQ treatment enhanced survival rate of physiologically aged mice (*p* < 0.05), but showed no significant effect in *Atm*-/- mice (Fig. 5i). Generally, these data demonstrate a lifespan-extending benefit of ATM activation by CQ.

**Figure 5:**
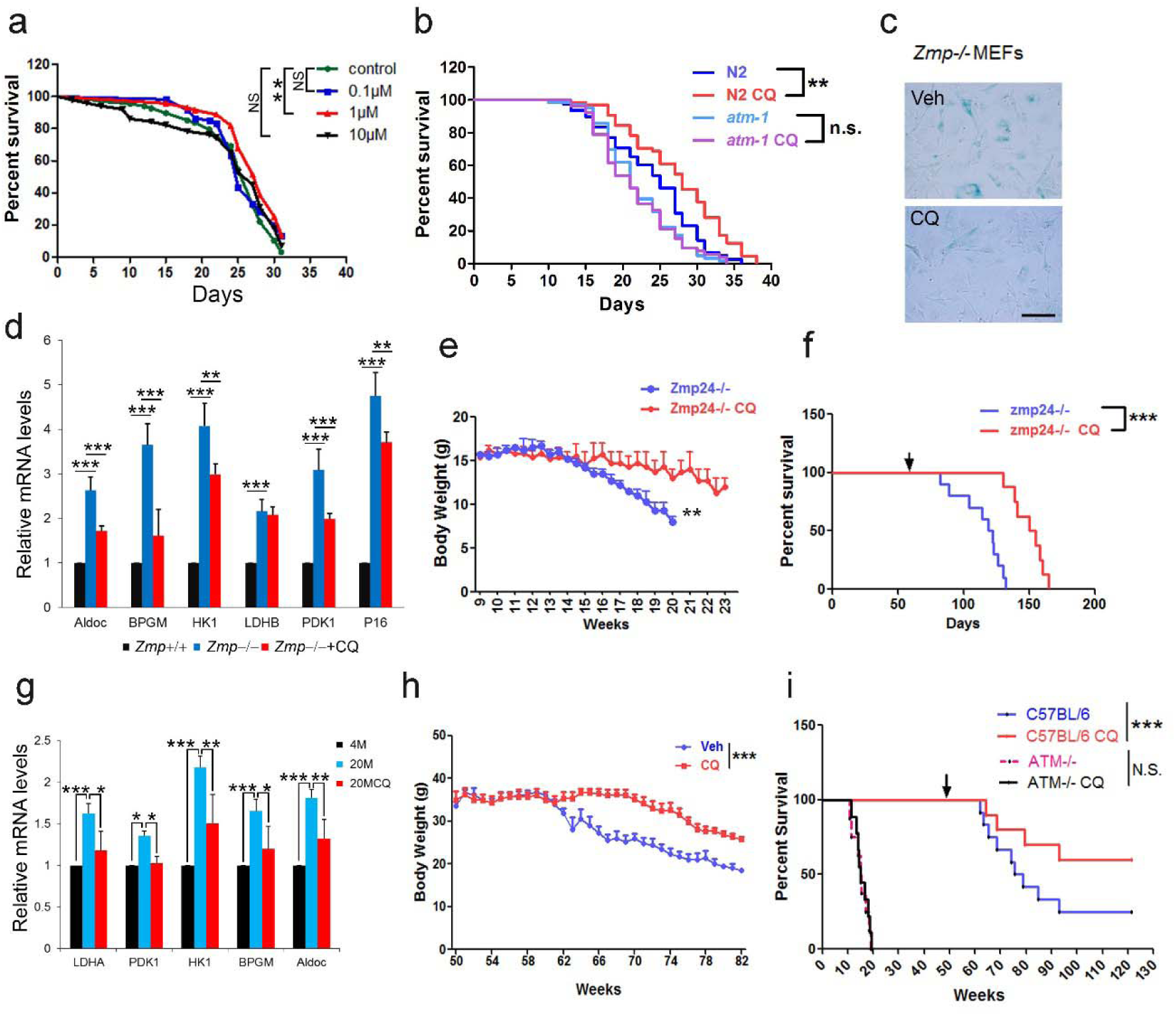
ATM Activation via CQ promotes longevity. (**a**) Survival analysis of *C. elegans* treated with the indicated dosage of CQ. \*\**P* < 0.01. NS indicates no significant difference. (**b**) Survival analysis of wild-type and *atm-1* null *C. elegans* cultured in medium with or without 1µM CQ. (**c**) Representative images showing SA-β-Gal staining in *Zmpste24*-/- MEFs with or without CQ treatment. Scale bar, 100 µm. (**d**) Quantitative RT-PCR analysis of mRNA levels of p16^Ink4a^ and indicated glycolytic genes in liver tissues of *Zmpste24*+/+, saline-treated, and CQ-treated *Zmpste24*-/- mice. Mice were treated for 8 weeks with two weekly intraperitoneal injections of CQ at 3.5 mg/kg. Data represent mean ± SEM. \**P* < 0.05, \*\**P* < 0.01, \*\*\**P* < 0.001. (**e**) Body weight of saline- and CQ-treated male *Zmpste24*-/- mice. Data represent mean ± SEM. \*\**P* < 0.01. (**f**) Kaplan-Meier survival curves of saline-treated (n = 12) and CQ-treated (n = 7) *Zmpste24*-/- mice. \*\*\**P* < 0.001. (**g**) Quantitative RT-PCR analysis of mRNA levels of indicated glycolytic genes in the liver tissues of 4-month-old, saline-treated 12-month-old (n = 3), and CQ-treated 20-month-old (n = 3) mice. Mice were treated for 8 weeks with two weekly intraperitoneal injections of CQ at 3.5 mg/kg. Data represent mean ± SEM. \**P* < 0.05, \*\**P* < 0.01, \*\*\**P* < 0.001. (**h**) Body weight of saline- and CQ-treated male aging mice. Data represent mean ± SEM. \*\*\**P* < 0.001. (**i**) Kaplan-Meier survival curves of saline-treated and CQ-treated aging C57BL/6J mice (n = 12 for each group) and *Atm*-/- mice (n = 9 for each group). \**P* < 0.05.

## Discussion

DNA damage accumulates with age and defective DNA repair accelerates aging. However, whether boosting DNA repair machinery promotes healthiness and longevity is still obscure. DNA damage stimulates DDR, but if persisted, it rather leads to senescence. Therefore, if enhancing DDR efficacy possibly promotes longevity, it must be DNA damage free. The antimalarial drug CQ could intercalate into the internucleosomal regions of chromatin, unwind DNA helical twist, and thus activate ATM without causing any DNA damage^6,54^. We demonstrate that long-term treatment with CQ activated ATM, slowed down senescence, restored age-related metabolic shift, and extended lifespan in both nematodes and mice. Mechanistically, ATM phosphorylated longevity gene SIRT6^55^, and prevented the MDM2-mediated ubiquitination and proteasomal degradation of SIRT6. To our knowledge, it is the first time establishing direct causal links between robust DNA repair machinery and longevity. Supporting this notion, the DNA repair efficacy is found enhanced in long-lived naked mole rat^56^, and human longevity is associated with single nucleotide polymorphisms (SNPs) in DNA repair genes/pathways^57,58^. Specifically, an *ATM* SNP that could enhance the transcription of *ATM* is associated with longevity in both Chinese and Italian populations^59,60^. In future study, it would be worthwhile to evaluate whether extra copies of *Atm* could promote longevity in model organisms.

Accumulation of DNA damages and metabolic disturbance are common denominators of aging^23,61^. Metabolic reprogramming from TCA cycle to glycolysis is prominent in both physiological and pathological aging^24,62^. Why senescent cells become glycolytic is poorly understood. Although it is recognized that deficiency in DNA repair machinery like ATM, WRN and Ercc1 accelerates aging and causes severe metabolic disorders^63,64^, it is unclear whether overexpression or enhancing DNA repair capacity, e.g. activating ATM, could not only enhance genomic stability but also hold off age-onset metabolic reprogramming and even extend lifespan. In our study, depletion of *ATM* induced metabolic reprogramming to glycolysis in senescent cells and premature ageing mouse models. Activation ATM by CQ reduced DNA damages and inhibited glycolysis. Remarkably, the long-term treatment of CQ extended lifespan in nematodes and mice. The data suggest a pro-longevity role of CQ-activated ATM, not only in improvement of DNA repair capacity, but also in the inhibition of glycolysis. Further, SIRT6 with diverge functions in biological process including DNA repair and metabolism, was discovered as the novel and direct target of ATM.

Recently, Bohr’s group uncovered an increased consumption of NAD+ by an early DDR factor poly (ADP-ribose) polymerase (PARP1), owing to accumulated DNA damages, accelerates aging in *Atm* mutant mice^65^. NAD^+^ serves as a cofactor of sirtuins, including SIRT1 and SIRT6. Therefore this work establishes a linear causal link between deficient DDR, DNA damage accumulation, consumption of NAD+, decline in sirtuin activity and aging. Moreover, administration of nicotinamide mononucleotide or nicotinamide riboside ameliorates age-related function decline and extends lifespan in mice^66,67^. Here, we found that *ATM* decline during aging caused DNA damage accumulation and enhanced glycolysis, both of which consume most of NAD^+^, providing an explanation of low NAD^+^ level in aged and *Atm*-/- mice.

Closely resembling normal ageing, HGPS has attracted numerous efforts in understanding of molecular mechanisms and developing therapeutic strategies^68,69^. We and others have found HGPS and *Zmpste24* null cells undergo accelerated senescence owing to defective chromatin remodeling^53,70–72^, delayed DDR and impaired DNA repair^12,72,73^. Specifically, Atm-Kap-1 signaling is compromised and SIRT6 protein and deacetylase activity are reduced in progeria cells^70,74^. Here we showed that Atm was significantly downregulated, which well explains the reduced SIRT6, delayed DDR and metabolic shift in progeria cells and mice. It would be interesting to investigate whether ectopic *Atm* or *Sirt6* could rescue progeroid features in these mice. Nevertheless, the pharmaceutical activation of ATM via CQ remarkably improved glucose homeostasis, DNA damage clearance, increased running endurance, and extended lifespan in progeria mice.

CQ is FDA-approved and clinically used medicine for treatment of malaria^75^. Through the activation of ATM, long-term treatment of CQ protects against atherosclerosis, improves insulin sensitivity, and rescues glucose tolerance in type 2 diabetes (T2D) models^76–78^. Lysosomotropic property of CQ also make it as a potent inhibitor of autophagy^79^. The dose (7.0 mg/kg/week in mice) applied in this study is equivalent to 35 mg/week for humans, much lower than that suggested for antimalarial treatment (500 mg/week, maximum 0.8 µM in plasma) and that for cancer therapy via autophagic inhibition (100–500 mg/day)^80^. Indeed, no obvious autophagic inhibition was observed in our study. The administration of CQ was conducted by intraperitoneal injection as described^78^. It seems that the frequently induced stress by repeated intraperitoneal injection imposes negative effect on lifespan, as saline-treated animals died earlier than untreated. Nonetheless, we treated the two groups of animals under the same conditions (age, diet, health status), and found that CQ-treated C57BL/6J mice exhibited much longer lifespan than saline-treated. When we summarized the lifespan data, i.e. ~840 days after birth of tested animals, which is equivalent to the median lifespan of C57BL/6J^35,67,81^, more than 60% CQ-treated mice are still alive. Therefore, we believe that the effect of CQ is not just an adaption to chronic stress induced by intraperitoneal injection but rather reflects a general regulation of murine lifespan.

In conclusion, our data establish direct causal links between robust DNA repair machinery and longevity. Together with DNA damage theory of aging, we propose that DNA damage activates DDR, however its constant activation causes senescence; defective ATM-SIRT6 axis underlies premature ageing, exemplified by HGPS and A-T mouse models, which are rescued by treatment of CQ and *Sirt6* transgene respectively; in physiological ageing, DNA damage-free activation of ATM by CQ stabilizes SIRT6, thus promoting longevity in both nematodes and mice (Supplementary Fig 8). Our findings also provide a novel therapeutic strategy for HGPS, and could facilitate clinical trials of CQ as an effective treatment for aging-related diseases.

## Experimental Procedures

### Mice

*Zmpste24*-/- mice and *Atm*-/- mice have been described previously^20,52^. *Sirt6*-transgenic mice (*Sirt6*-tg) in C57BL/6J background were constructed by injecting cloned mSirt6 cDNA with CAG promoter into fertilized eggs. Primers for genotyping of *Sirt6* transgenic allele were as follows; forward: 5’-CTGGTTATTGTGCTGTCTCATCAT-3’ and reverse: 5’-CCGTCTACGTTCTGGCTGAC-3’. *Atm*-/- mice were crossed to *Sirt6*-tg mice to get *Atm-/-; Sirt6*-tg mice. Chloroquine (CQ) experiments were conducted as described^78^. Briefly, 12-month-old wild-type C57BL/6J male mice and 2-month-old *Zmpste24*-/- male mice were administered with CQ (Sigma, St. Louis, MO) in 0.9% saline twice per week at 7 mg/kg body weight, and control group was treated with saline alone. 8 weeks after treatment of CQ, mice were subjected for functional tests. Body weight and lifespan was recorded. The survival rate was analyzed by Kaplan–Meier method and statistical comparison was performed by Log-rank Test. Mice were housed and handled in the laboratory animal research center of Shenzhen University. All experiments were performed in accordance with the guidelines of the Institutional Animal Care and Use Committee (IACUC).

### *C. elegans* survival assay

*C. elegans* nematode survival assay was performed according to standard protocols^82^. Briefly, wild-type and *atm-1* null nematodes (100 to 150 per group) synchronized to prefertile young adult stage were exposed to NGM plates containing the indicated dosage of CQ. After 1-day incubation, animals were transferred to fresh incubation plates without CQ for another 2 days. This procedure was repeated every three days. Nematodes that showed no response to gentle stimulation were recorded as dead. The survival data was analyzed by Kaplan–Meier method and statistical comparison was performed by Log-rank Test.

### Cell lines

HEK293, immortalized mouse embryonic fibroblasts (MEFs), human skin fibroblasts (HSFs), HepG2 and U2OS cells were cultured in Gibco^®^ DMEM (Life Technologies, USA) with 10% fetal bovine serum (FBS), 100 U/ml penicillin and streptomycin (P/S) at 37 °C in a 5% CO_2_ humidified atmosphere. Primary MEFs were generated from 13.5-day-old embryos as described^12^, and were cultured in DMEM supplemented with 0.1 mM nonessential amino acids (Life Technologies), 0.1 mM 2-mercaptoethanol (Sigma-Aldrich, USA). For CQ experiments, cells were maintained in the medium containing 1μM chloroquine for 12 h, and then grown in new fresh medium for 48 h.

| Cell lines | Description and Application in this study |
| --- | --- |
| HEK293 | Transfection; Immunoprecipitation |
| ATM KO HEK293 | Constructed by CRISPR-cas9 system; Ubiquitination and Degradation; p-S/T Q level of ectopic SIRT6 |
| WT, *Atm*<sup>-/-</sup> MEFs and *Zmpste24*<sup>-/-</sup> MEFs | Prepared from WT, *Atm*<sup>-/-</sup> and *Zmpste24*<sup>-/-</sup> embryonic mice; Senescence assay; Glycolysis |
| WT and *Atm*<sup>-/-</sup> MEFs | RNA-seq; Metabolome; Degradation assay of endogenous Sirt6 protein |
| HSF cells | shATM by lentivirus; Senescence assay; Glycolysis |
| HepG2 cells | siSIRT6/siATM to perform the metabolic assay |
| U2OS cells | DNA repair; Endogenous SIRT6 protein level |

### Plasmids

Human Flag-SIRT6, pcDNA3.1 Flag-ATM, Flag-ATM S1981A, and pcDNA3 human MDM2 were all purchased from Addgene (Cambridge, MA). Flag-SIRT6 with amino acid substitution mutations (S112A, S112D, K346R/K349R) were generated by PCR-based mutagenesis using pcDNA3-Flag-SIRT6 as a template and QuikChange II site-directed mutagenesis kit (Agilent Technologies), following the manufacturer’s instruction. Primer sequences for amino acid mutations of SIRT6 were as follows: SIRT6 S112A: (forward) 5’-cgtccacgttctgggcgaccaggaagcgga-3’, (reverse) 5’-tccgcttcctggtcgcccagaacgtggacg-3’; SIRT6 S112D: (forward) 5’-ccgtccacgttctggtcgaccaggaagcggag-3’, (reverse) 5’-ctccgcttcctggtcgaccagaacgtggacgg-3’; SIRT6 K346R: (forward) 5’-ggccttcacccttctggggggtctgtg-3’, (reverse) 5’-cacagaccccccagaagggtgaaggcc-3’; SIRT6 K349R: (forward) 5’-gccttggccctcacccttttggggggt-3’, (reverse) 5’-accccccaaaagggtgagggccaaggc-3. HA-tagged human SIRT6 plasmid was amplified from their respective cDNAs and constructed into pKH3-HA vector. To express four truncated forms of SIRT6 protein, HA-SIRT6 plasmid as a template was constructed by PCR-based deletion.

### Protein extraction and Western blotting

For whole cell protein extraction, cells were suspended in 5 volumes of suspension buffer (20 mM Tris-HCl, pH 7.5, 150 mM NaCl, 1 mM EDTA, 1 mM DTT, protease inhibitor cocktail), and then added 5 volumes of 2X SDS loading buffer and incubated at 98 °C for 6 min. Mice tissues were homogenized with 1mL of ice-cold tissue lysis buffer (25 mM TrisHCl, pH 7.5, 10 mM Na_3_VO_4_, 100 mM NaF, 50 mM Na_4_P_2_O_7_, 5 mM EGTA, 5 mM EDTA, 0.5% SDS, 1% NP-40, protease inhibitor cocktail). After homogenization and sonication, lysates were centrifuged at 16,000 g for 15 min. The clean supernatant was carefully transferred to new tubes. Protein concentrations were determined by using bicinchoninic acid (BCA) assay method (Pierce, Rockford, IL) and were normalized with lysis buffer for each sample. Samples were denatured in 1X SDS loading buffer by boiling at 98 °C for 6 min. Proteins were separated by loading to SDS-polyacrylamide gels, and then were transferred to PVDF membrane (Millipore). The protein levels were determined by immunoblotting using respective antibodies. The ImageJ program was used for densitometric analysis of immunoblotting, and the quantification results were normalized to the loading control.

### Antibodies

Rabbit anti-SIRT6 (ab62739), ATM (ab78), SIRT1 (ab12193), γH2AX (ab81299), RAP80 (ab52893), Kap-1 (ab10484), p-KAP-1 (Ser824, ab70369), PDPK1 (ab52893) antibodies were obtained from Abcam (Cambridge, UK). Anti-lamin A/C (sc-20681), p21 (sc-6246), MDM2 (sc-965), P53 (sc-6243) antibodies were purchased from Santa Cruz Biotechnology. Rabbit anti-γH2AX (05–636), p-ATM (Ser1981) (05–740), histone H3 (07–690), anti-H3K56ac (07–677) and H3K9ac (07–352) antibodies were sourced from EMD Millipore. Mouse anti-p-ATM (Ser1981) (#5883), p-S/TQ (#9607), Ubiquitin (#3936), AMPK (#5831), rabbit anti-p-Akt (Ser308) (#9275), p-Akt (Ser473) (#9271), total Akt (#9272), p-S6 (#2211), S6 (#2217), p-GSK3β (Ser9) (#9322), GSK3β (#5676), p-AMPK (Thr172) (#2535), cleaved caspase-3 (#9661) antibodies were purchased from Cell Signaling Technology (Beverly, MA). Mouse anti- HA, Flag, rabbit anti-LC3B, P62 antibodies were obtained from Sigma-Aldrich. Anti-Nbs1 (NB100–143) antibody was purchased from Novus Biologicals. Mouse anti-actin, tubulin antibodies were obtained from Beyotime. Anti-pS112 SIRT6 monoclonal antibodies were prepared by Abmart generating from a specific phosphorylated peptide (peptide sequence CLRFVSPQNV).

### Protein degradation assay

HEK293 cells (WT and ATM-deficient cells) were transfected with Flag-SIRT6 alone or together with Mdm2. 48 h later, the cells were treated with 50 μg/ml of cycloheximide (CHX, Sigma-Aldrich), a translation inhibitor. For endogenous SIRT6 protein degradation assay, ATM wild-type and null MEFs were grown in 6-cm plates, and were treated with 50 mg/ml CHX for indicated time points. Cells were collected and the protein levels were determined by western blotting, the subsequent quantification was performed with ImageJ^®^ software.

### *In vivo* ubiquitination assay

*In vivo* ubiquitination assay was performed by transfecting HEK293 cells in 6-cm dishes with 1 μg Myc-ubiquitin, 2 μg Flag-SIRT6 or its mutations, and/or 1 μg MDM2 vector. 48 h after transfection, cells were lysed in the buffer (25 mM Tris-HCl pH 8.0, 250 mM NaCl, 10 mM Na_3_VO_4_, 1mM EDTA, 10% glycerol, protease inhibitor cocktail, and 0.1 mM Phenylmethylsulphonyl fluoride), and then incubated with Flag-M2 beads (Sigma-Aldrich) overnight at 4 °C. Beads were washed with lysis buffer for three times, bound proteins were eluted by adding 1.5 × SDS loading buffer. The ubiquitin levels were analyzed by immunoblotting.

### *In vitro* kinase assay

HEK293T cells were transfected with 10 μg of FLAG-ATM and then treated with CPT. Activated ATM was immune-purified from the cell extracts with FLAG beads (Sigma, M8823). GST-SIRT6 or the S112A mutant was purified from bacteria. Kinase reactions were initiated by incubating purified ATM with GST-SIRT6 in the kinase buffer with or without 1mM ATP for 120 min at 30□. After reaction, proteins were blocked by SDS loading buffer. The membrane was then subjected to Western blotting with antibodies against p-S/TQ.

### Immunoprecipitation

Cells under indicated treatments were totally lysed in lysis buffer containing 20 mM HEPES, pH 7.5, 150 mM NaCl, 10 mM Na_3_VO_4_, 10% glycerol, 2 mM EDTA, protease inhibitor cocktail, and 0.1 mM Phenylmethylsulphonyl fluoride. After sonication and centrifugation, the supernatant was collected and incubated with H3K9ac (Millipore, 2 μg/sample) overnight at 4 □ with a gentle rotation. Protein A/G agarose (Pierce, 10 μl/sample) were added to the tubes and rotated at 4 □ for 2 h. Beads were precipitated by centrifugation at 1000 g for 15 s and washed three times with cold lysis buffer. The pellet was resuspended in 1.5 × SDS loading buffer and incubated at 98 □ for 6 min. The supernatants were collected and used for western blotting.

### GST pull-down assay

A series of GST fusion proteins of truncated ATM, which together spanned the full-length of ATM, were constructed into pGEX4T-3 vector. For GST pull-down, bacterially expressed 6 × His-tagged SIRT6 was separately incubated with various GST-ATM fragments in a buffer of 150 mM NaCl, 20 mM Tris-HCl [pH 7.5], 5 mM MgCl2, 0.2 mM EDTA, 10% glycerol, 0.2% NP-40 and protease inhibitors (Roche Complete). GST-fusion proteins were then precipitated by adding Glutathione Sepharose^®^ fast flow (GE Healthcare). After washed twice with TEN buffer (0.5% Nonidet P-40, 20 mM Tris-HCl [pH 7.4], 0.1 mM EDTA, and 300 mM NaCl), glutathione agarose beads were analyzed by Western blotting and coomassie staining.

### RNA interference

Briefly, cells were transfected with the indicated small interfering RNAs (siRNAs) for 48 h using Lipofectamine 3000 (Invitrogen) according to the manufacturer’s instructions. The following siRNA oligonucleotides targeting human ATM, SIRT6 and HDM2 were purchased from Shanghai GenePharma, China. Si-*ATM* (AAUGUCUUUGAGUAGUAUGUU^83^, #2: AAGCACCAGUCCAGUAUUGGC^84^); si-SIRT6 (AAGAAUGUGCCAAGUGUAAGA); si*-HDM2* (#1: AACGCCACAAATCTGATAGTA; #2: AATGCCTCAATTCACATAGAT). A scrambled siRNA sequence was used as control. Lentiviral shRNA constructs were generated in a pGLVH1 backbone (GenePharma, China), and virus was produced in HEK293 cells. To deplete ATM in HSF cells, lentiviral infection was performed in the presence of 5 μg/ml polybrene. 2 days later, the infected HSF cells were selected with 2 μg/ml puromycin. To downregulate sir-2.4 expression, the NL2099 worms were exposed to incubation plates containing HT115 bacteria with sir-2.4 RNAi vector.

### CRISPR/Cas9-mediated genome editing

Gene mutagenesis by CRISPR/Cas9 system was conducted as described^85^. The following gRNA oligonucleotides targeting human ATM, SIRT6 were constructed in pX459 vector (Addgene, #48139). sg*ATM* F: 5’- CACCGATATGTGTTACGATGCCTTA -3′, R: 5′-AAACTAAGGCATCGTAACACATATC -3′. HEK293 cells were transfected with pX459 or pX459-gRNA by using Lipofetamine^®^ 3000 Transfection Reagent according to manufacturer introductions. After 2-day culture, cells were selected with 2 μg/ml puromycin, six colonies were picked and grown to establish stable cell lines. The targeted mutations were identified by Western blotting, and PCR-based sequencing.

### EdU (5-ethynyl-2’-deoxyuridine) incorporation assay

EdU incorporation assays were conducted in HSF cells to estimate cell proliferation using the Click-iT™ EdU Alexa Fluor^®^ 488 Kit (Invitrogen). HSF cells, infected by the respective lentiviruses containing shNC and sh*ATM*, were cultured in a 6-well plates containing the coverslips in the presence of 10μM EdU for 12 hours. Cells were fixed in 3.7 % formaldehyde followed by a 0.5% Triton X-100 permeabilization, and then stained with Alexa Fluor picolyl azide. Five random views were captured to calculate the positive staining rate for each group.

### Growth curves and SA-β-gal assays

Cell population doublings were monitored using a Coulter Counter. SA-β-galactosidase assay in primary cells was performed using Senescence beta-galactosidase staining Kit (#9860, CST) according to manufacturer instructions, five views were captured randomly to calculate the positive staining rate for each group.

### RNA Preparation and Real-Time qPCR

Total RNA was extracted from cells or the tissues of mice using Trizol^®^ reagent RNAiso Plus (TaKaRa, Japan) following the phenol–chloroform extraction method. Purified total RNA was used to obtain cDNA using PrimeScript^TM^ RT Master Mix (Takara, Japan) following this method: 37 °C for 30 min, and 85 °C for 5 s. The gene expression was analyzed with CFX Connected^TM^ Real-Time PCR Detection System (BioRad) with SYBR^®^ Ex Taq Premixes (Takara, Japan). Gene expression levels were normalized to actin.

### Glucose tolerance test

Mice were fasted overnight (6 p.m. to 9 a.m.), and D-glucose (2.5 g/kg body weight) was administrated intraperitoneally. Blood glucose levels were determined from tail vein blood using glucometer (Onetouch^®^ ultravue, Johnson) at 0, 30, 60, 90, and 120 min after D-glucose injection.

### Insulin tolerance test

Mice were fasted for 6 hours (8 a.m. to 2 p.m.), and recombinant human insulin (0.75 U/kg body weight) was administered intraperitoneally. Blood glucose levels were determined from tail vein blood using glucometer (Onetouch^®^ ultravue, Johnson) at 0, 30, 60, 90, and 120 min after insulin injection.

### Lactate assay

Mouse serum was five-fold diluted, and lactate concentration was determined with the Lactate Colorimetric Assay Kit (BioVision).

### Endurance running test

*Zmpste24*-/- mice were treated for 8 weeks with chloroquine or saline before running on a Rota-Rod Treadmill (YLS-4C, Jinan Yiyan Scientific Research Company, Shandong, China) to test the effect of chloroquine on fatigue resistance. Mice were placed on the rotating lane, and the speed was gradually increased to 10 r/min. When mice were exhausted and safely dropped from the rotating lane, the time latency to fall were automatically recorded.

### Metabolite analysis

Wild-type and *ATM* KO cells were grown in normal medium for 24 hours, and methanol-fixed cell pellets were analyzed by two liquid chromatography-tandem mass spectrometry (LC-MS) method as described^86^.

### Immunofluorescence microscopy

The cells were fixed using 4% paraformaldehyde at room temperature for 15 min, and then permeabilized using 0.5% Triton X-100 at room temperature for 10 min, blocked using 10% FBS/PBS, and stained using primary antibodies diluted in PBS containing 2% BSA overnight at 4 □. The primary antibodies were detected using an Alexa-488-conjugated anti-mouse secondary antibody (Invitrogen). The nuclei were stained using DAPI in anti-fade mounting medium. Images were captured using Zeiss LSM880 confocal/multiphoton microscope.

### ChIP assay

Cells were fixed in 1% formaldehyde for 10 min at room temperature. The crosslinking reaction was quenched with 0.125 M glycine. After washed with PBS, cells were lysed with lysis buffer (50mM Tris·HCl pH 8.0, 2mM EDTA, 15 mM NaCl, 1% SDS, 0.5% deoxycholate, protease inhibitor cocktail, and 1mM PMSF). After sonication and centrifugation, the supernatant was collected and precleared in dilution buffer (50mM Tris-HCl pH 8.0, mM EDTA, 150mM NaCl, 1% Triton X-100) with protein A/G Sepharose and pre-treated salmon DNA. The precleared samples were incubated overnight with H3K9ac antibody (2 μg/sample, Millipore) or appropriate control IgGs (Santa Cruz), and protein A/G Sepharose (Invitrogen). After washed sequentially with a series of buffers, the beads were heated at 65 °C to reverse the cross-link. DNA fragments were purified and analyzed. Real-time PCR was performed with primers as described^31^:

LDHB-ChIP-5′: AGAGAGAGCGCTTCGCATAG LDHB-ChIP-3′: GGCTGGATGAGACAAAGAGC ALDOC-ChIP-5′: AAGTGGGGCACTGTTAGGTG ALDOC-ChIP-3′: GTTGGGGATTAAGCCTGGTT PFKM-ChIP-5′: TTAAGACAAAGCCTGGCACA PFKM-ChIP-3′: CAACCACAGCAATTGACCAC LDHA-ChIP-5′: AGGGGGTGTGTGAAAACAAG LDHA-ChIP-3′: ATGGCTTGCCAGCTTACATC LDHA-ChIP-1Kb-5′: TGCAAGACAAGTGTCCCTGT LDHA-ChIP-1Kb-3′: GAGGGAATGAAGCTCACAGC

### Statistical analysis

Statistical analyses were conducted using two-tailed Student’s t-test between two groups. All data are presented as Mean ± S.D. or Mean ± S.E.M. as indicated, and a *P* value < 0.05 was considered statistically significant.

### Primers for q-PCR

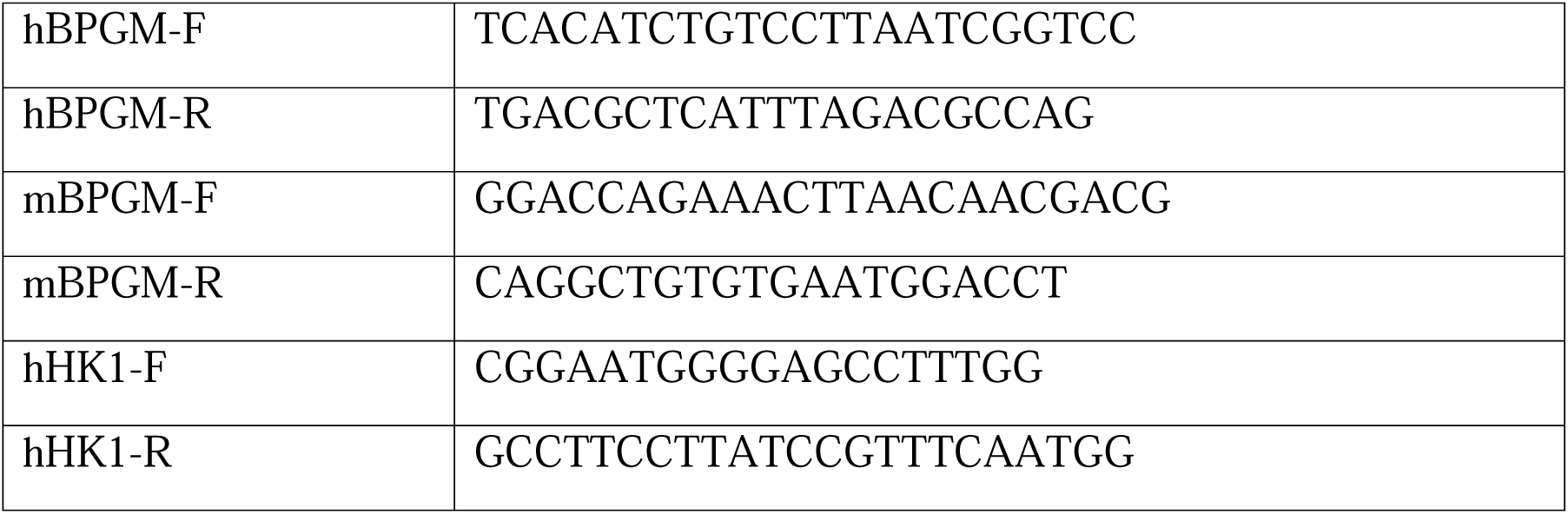

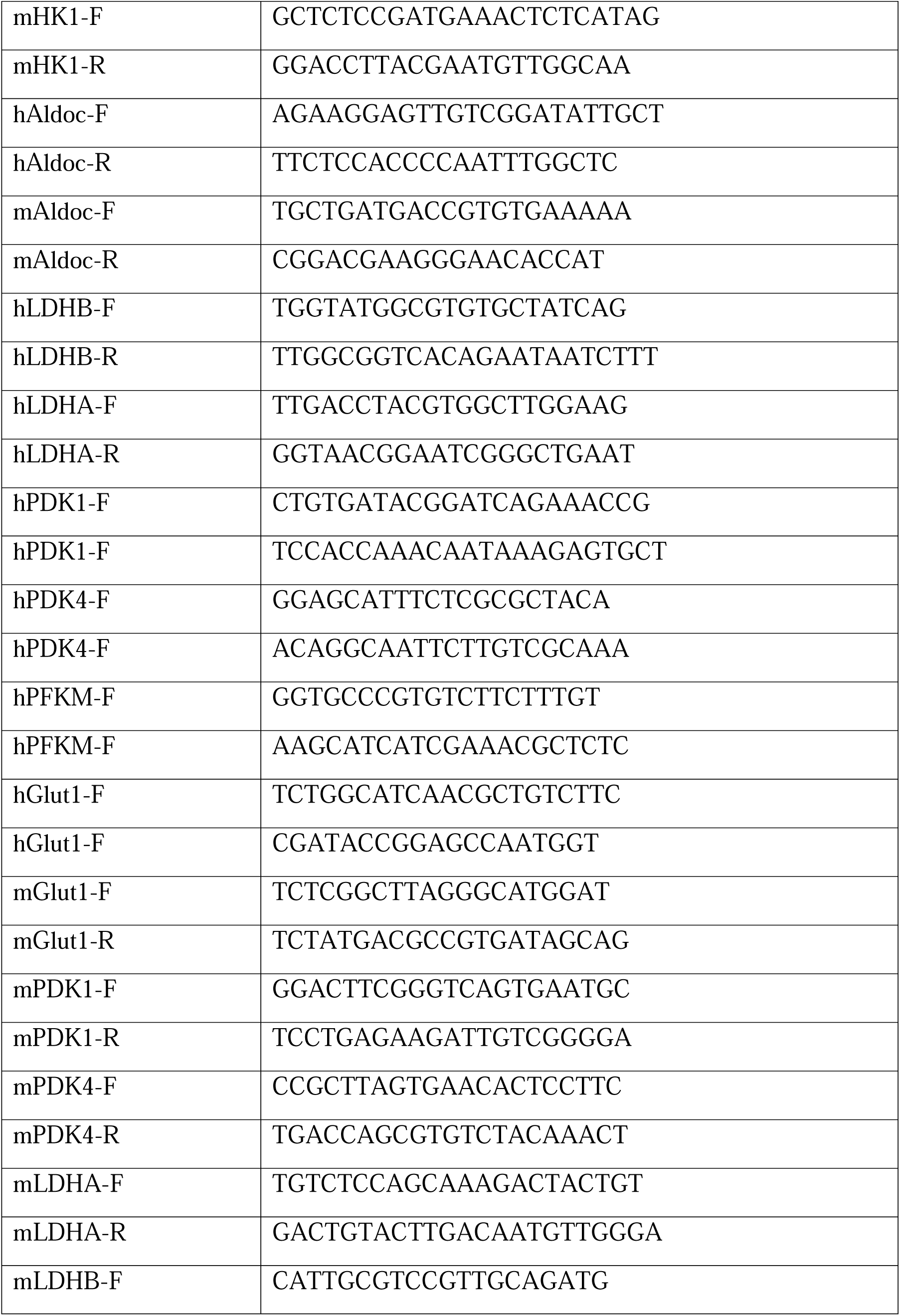

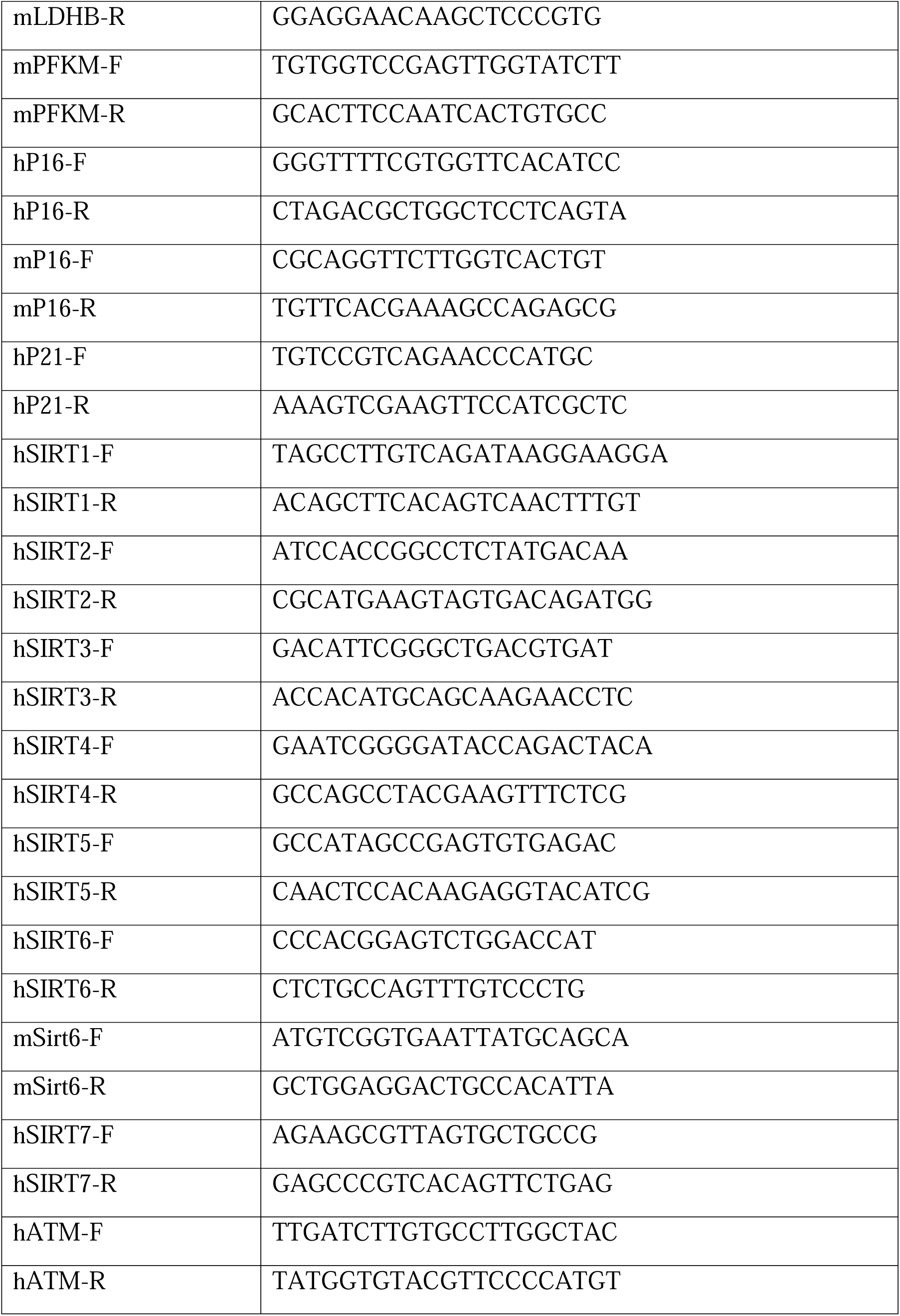

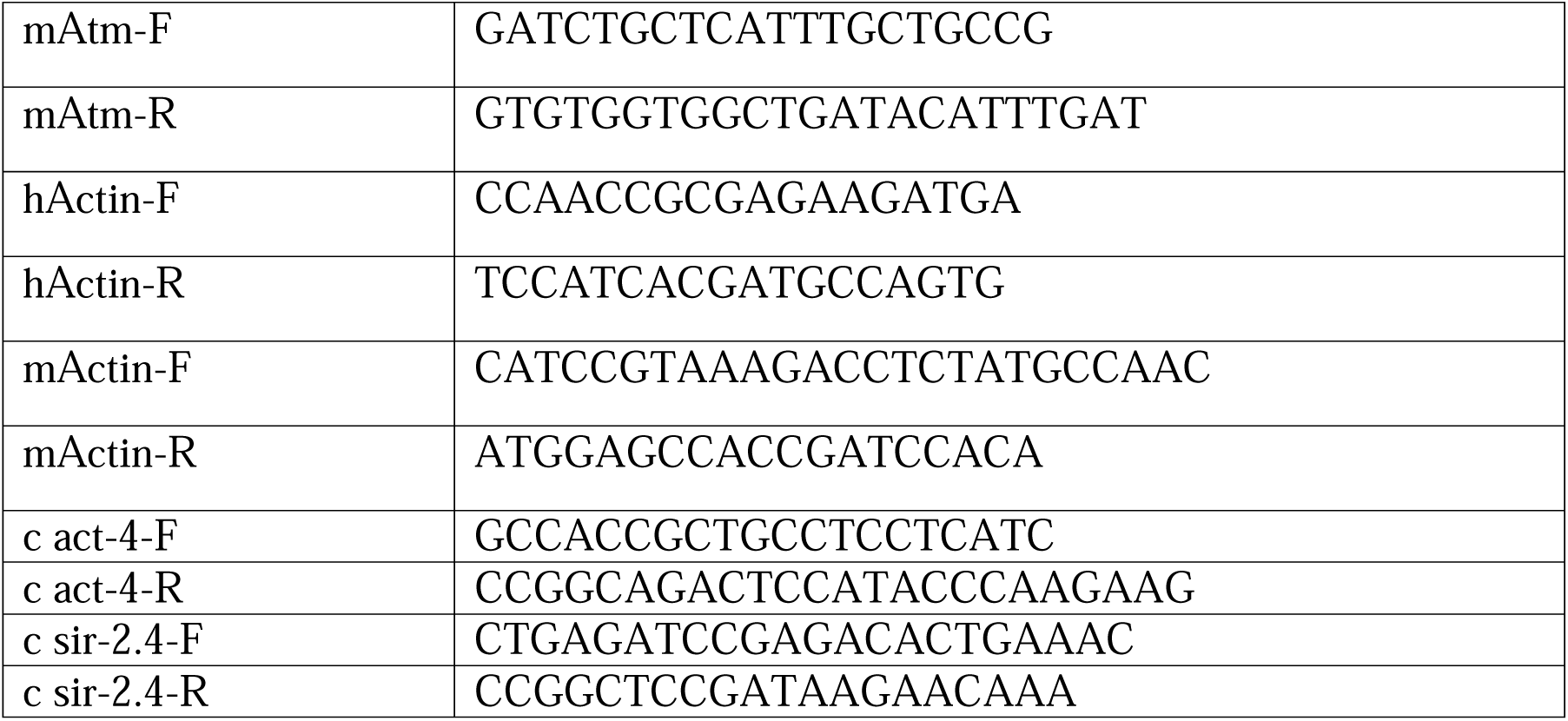

## Acknowledgements

We thank Dr Yosef Shiloh (Tel Aviv University, Israel) for *Atm-/-; p53*-/- MEF cells and Dr. Raul Mostoslavsky (Massachusetts General Hospital Cancer center, USA) for *Sirt6*-/- MEFs. This project is supported by research grants from National Natural Science Foundation of China (81422016, 91439133 and 81571374), Natural Science Foundation of Guangdong Province (2014A030308011, 2015A030308007), Shenzhen Peacock Plan (KQTD20140630100746562) and Shenzhen Science and Technology Innovation Commission (CXZZ20140903103747568).

## Reference

1 Lombard, D. B. et al. DNA repair, genome stability, and aging. Cell 120, 497–512, doi:10.1016/j.cell.2005.01.028 (2005).

2 Vijg, J. Somatic mutations, genome mosaicism, cancer and aging. Current opinion in genetics & development 26, 141–149, doi:10.1016/j.gde.2014.04.002 (2014).

3 Halliwell, B. & Whiteman, M. Measuring reactive species and oxidative damage in vivo and in cell culture: how should you do it and what do the results mean? Br J Pharmacol 142, 231–255 (2004).

4 Tanaka, T., Halicka, H. D., Huang, X., Traganos, F. & Darzynkiewicz, Z. Constitutive histone H2AX phosphorylation and ATM activation, the reporters of DNA damage by endogenous oxidants. Cell Cycle 5, 1940–1945 (2006).

5 Guleria, A. & Chandna, S. ATM kinase: Much more than a DNA damage responsive protein. DNA repair 39, 1–20, doi:10.1016/j.dnarep.2015.12.009 (2016).

6 Bakkenist, C. J., & Kastan, M. B. DNA damage activates ATM through intermolecular autophosphorylation and dimer dissociation. Nature 421, 499–506, doi:10.1038/nature01368 (2003).

7 Paull, T. T. Mechanisms of ATM Activation. Annual review of biochemistry 84, 711–738, doi:10.1146/annurev-biochem-060614-034335 (2015).

8 Burma, S., Chen, B. P., Murphy, M., Kurimasa, A. & Chen, D. J. ATM phosphorylates histone H2AX in response to DNA double-strand breaks. J Biol Chem 276, 42462–42467, doi:10.1074/jbc.C100466200 (2001).

9 Maslov, A. Y., & Vijg, J. Genome instability, cancer and aging. Biochim Biophys Acta 1790, 963–969, doi:10.1016/j.bbagen.2009.03.020 (2009).

10 Harper, J. W. & Elledge, S. J. The DNA damage response: ten years after. Mol Cell 28, 739–745, doi:S1097-2765(07)00783-6 [pii] 10.1016/j.molcel.2007.11.015 (2007).

11 Li, Z. et al. Impaired DNA double-strand break repair contributes to the age-associated rise of genomic instability in humans. Cell death and differentiation, doi:10.1038/cdd.2016.65 (2016).

12 Liu, B. et al. Genomic instability in laminopathy-based premature aging. Nature medicine 11, 780–785, doi:10.1038/nm1266 (2005).

13 Hoeijmakers, J. H. DNA damage, aging, and cancer. The New England journal of medicine 361, 1475–1485, doi:10.1056/NEJMra0804615 (2009).

14 Shiloh, Y. & Lederman, H. M. Ataxia-telangiectasia (A-T): An emerging dimension of premature ageing. Ageing research reviews, doi:10.1016/j.arr.2016.05.002 (2016).

15 Boder, E. & Sedgwick, R. P. Ataxia-telangiectasia; a familial syndrome of progressive cerebellar ataxia, oculocutaneous telangiectasia and frequent pulmonary infection. Pediatrics 21, 526–554 (1958).

16 Lebel, M., Spillare, E. A., Harris, C. C. & Leder, P. The Werner syndrome gene product co-purifies with the DNA replication complex and interacts with PCNA and topoisomerase I. J Biol Chem 274, 37795–37799 (1999).

17 Balajee, A. S. et al. The Werner syndrome protein is involved in RNA polymerase II transcription. Mol Biol Cell 10, 2655–2668 (1999).

18 Cooper, M. P. et al. Ku complex interacts with and stimulates the Werner protein. Genes Dev 14, 907–912 (2000).

19 Li, B. & Comai, L. Functional interaction between Ku and the werner syndrome protein in DNA end processing. J Biol Chem 275, 28349–28352 (2000).

20 Barlow, C. et al. Atm-deficient mice: a paradigm of ataxia telangiectasia. Cell 86, 159–171 (1996).

21 Hoeijmakers, J. H. DNA damage, aging, and cancer. N Engl J Med 361, 1475–1485, doi:10.1056/NEJMra0804615 (2009).

22 Hasty, P. The impact of DNA damage, genetic mutation and cellular responses on cancer prevention, longevity and aging: observations in humans and mice. Mech Ageing Dev 126, 71–77, doi:10.1016/j.mad.2004.09.036 (2005).

23 Lopez-Otin, C., Blasco, M. A., Partridge, L., Serrano, M. & Kroemer, G. The hallmarks of aging. Cell 153, 1194–1217, doi:10.1016/j.cell.2013.05.039 (2013).

24 Shimizu, I., Yoshida, Y., Suda, M. & Minamino, T. DNA damage response and metabolic disease. Cell Metab 20, 967–977, doi:10.1016/j.cmet.2014.10.008 (2014).

25 Lopez-Otin, C., Galluzzi, L., Freije, J. M., Madeo, F. & Kroemer, G. Metabolic Control of Longevity. Cell 166, 802–821, doi:10.1016/j.cell.2016.07.031 (2016).

26 Kruger, A. & Ralser, M. ATM is a redox sensor linking genome stability and carbon metabolism. Science signaling 4, pe17, doi:10.1126/scisignal.2001959 (2011).

27 Cosentino, C., Grieco, D. & Costanzo, V. ATM activates the pentose phosphate pathway promoting anti-oxidant defence and DNA repair. The EMBO journal 30, 546–555, doi:10.1038/emboj.2010.330 (2011).

28 Aird, K. M. et al. ATM couples replication stress and metabolic reprogramming during cellular senescence. Cell reports 11, 893–901, doi:10.1016/j.celrep.2015.04.014 (2015).

29 Denu, J. M. The Sir 2 family of protein deacetylases. Curr Opin Chem Biol 9, 431–440, doi:S1367-5931(05)00107-9 [pii] 10.1016/j.cbpa.2005.08.010 (2005).

30 Finkel, T., Deng, C. X. & Mostoslavsky, R. Recent progress in the biology and physiology of sirtuins. Nature 460, 587–591, doi:nature08197 [pii] 10.1038/nature08197 (2009).

31 Zhong, L. et al. The histone deacetylase Sirt6 regulates glucose homeostasis via Hif1alpha. Cell 140, 280–293, doi:10.1016/j.cell.2009.12.041 (2010).

32 Sebastian, C. et al. The histone deacetylase SIRT6 is a tumor suppressor that controls cancer metabolism. Cell 151, 1185–1199, doi:10.1016/j.cell.2012.10.047 (2012).

33 Mostoslavsky, R. et al. Genomic instability and aging-like phenotype in the absence of mammalian SIRT6. Cell 124, 315–329, doi:10.1016/j.cell.2005.11.044 (2006).

34 Mao, Z. et al. Sirtuin 6 (SIRT6) rescues the decline of homologous recombination repair during replicative senescence. Proc Natl Acad Sci U S A 109, 11800–11805, doi:10.1073/pnas.1200583109 (2012).

35 Kanfi, Y. et al. The sirtuin SIRT6 regulates lifespan in male mice. Nature 483, 218–221, doi:10.1038/nature10815 (2012).

36 Samper, E., Nicholls, D. G. & Melov, S. Mitochondrial oxidative stress causes chromosomal instability of mouse embryonic fibroblasts. Aging cell 2, 277–285 (2003).

37 Parrinello, S. et al. Oxygen sensitivity severely limits the replicative lifespan of murine fibroblasts. Nature cell biology 5, 741–747, doi:10.1038/ncb1024 (2003).

38 Sherr, C. J., & DePinho, R. A. Cellular senescence: mitotic clock or culture shock? Cell 102, 407–410 (2000).

39 Bar, R. S. et al. Extreme insulin resistance in ataxia telangiectasia: defect in affinity of insulin receptors. The New England journal of medicine 298, 1164–1171, doi:10.1056/NEJM197805252982103 (1978).

40 Espach, Y., Lochner, A., Strijdom, H. & Huisamen, B. ATM protein kinase signaling, type 2 diabetes and cardiovascular disease. Cardiovascular drugs and therapy 29, 51–58, doi:10.1007/s10557-015-6571-z (2015).

41 Hagen, T. M. et al. Mitochondrial decay in hepatocytes from old rats: membrane potential declines, heterogeneity and oxidants increase. Proc Natl Acad Sci U S A 94, 3064–3069 (1997).

42 Lenaz, G. et al. Mitochondrial bioenergetics in aging. Biochim Biophys Acta 1459, 397–404 (2000).

43 Kruiswijk, F., Labuschagne, C. F. & Vousden, K. H. p53 in survival, death and metabolic health: a lifeguard with a licence to kill. Nat Rev Mol Cell Biol 16, 393–405, doi:10.1038/nrm4007 (2015).

44 Schwartzenberg-Bar-Yoseph, F., Armoni, M. & Karnieli, E. The tumor suppressor p53 down-regulates glucose transporters GLUT1 and GLUT4 gene expression. Cancer Res 64, 2627–2633 (2004).

45 Mostoslavsky, R. et al. Genomic instability and aging-like phenotype in the absence of mammalian SIRT6. Cell 124, 315–329 (2006).

46 You, Z., Chahwan, C., Bailis, J., Hunter, T. & Russell, P. ATM activation and its recruitment to damaged DNA require binding to the C terminus of Nbs1. Molecular and cellular biology 25, 5363–5379, doi:10.1128/MCB.25.13.5363-5379.2005 (2005).

47 Assaily, W. et al. ROS-mediated p53 induction of Lpin1 regulates fatty acid oxidation in response to nutritional stress. Molecular cell 44, 491–501, doi:10.1016/j.molcel.2011.08.038 (2011).

48 Sarre, A., Gabrielli, J., Vial, G., Leverve, X. M. & Assimacopoulos-Jeannet, F. Reactive oxygen species are produced at low glucose and contribute to the activation of AMPK in insulin-secreting cells. Free Radic Biol Med 52, 142–150, doi:10.1016/j.freeradbiomed.2011.10.437 (2012).

49 Hickson, I. et al. Identification and characterization of a novel and specific inhibitor of the ataxia-telangiectasia mutated kinase ATM. Cancer research 64, 9152–9159, doi:10.1158/0008-5472.CAN-04-2727 (2004).

50 Berkovich, E., Monnat, R. J., Jr. & Kastan, M. B. Roles of ATM and NBS1 in chromatin structure modulation and DNA double-strand break repair. Nature cell biology 9, 683–690, doi:10.1038/ncb1599 (2007).

51 Thirumurthi, U. et al. MDM2-mediated degradation of SIRT6 phosphorylated by AKT1 promotes tumorigenesis and trastuzumab resistance in breast cancer. Sci Signal 7, ra71, doi:10.1126/scisignal.2005076 (2014).

52 Pendas, A. M. et al. Defective prelamin A processing and muscular and adipocyte alterations in Zmpste24 metalloproteinase-deficient mice. Nat Genet 31, 94–99, doi:10.1038/ng871 (2002).

53 Liu, B., Wang, Z., Ghosh, S. & Zhou, Z. Defective ATM-Kap-1-mediated chromatin remodeling impairs DNA repair and accelerates senescence in progeria mouse model. Aging Cell 12, 316–318, doi:10.1111/acel.12035 (2013).

54 Krajewski, W. A. Alterations in the internucleosomal DNA helical twist in chromatin of human erythroleukemia cells in vivo influences the chromatin higher-order folding. FEBS Lett 361, 149–152 (1995).

55 Tasselli, L., Zheng, W. & Chua, K. F. SIRT6: Novel Mechanisms and Links to Aging and Disease. Trends Endocrinol Metab, doi:10.1016/j.tem.2016.10.002 (2016).

56 MacRae, S. L. et al. DNA repair in species with extreme lifespan differences. Aging (Albany NY) 7, 1171–1184, doi:10.18632/aging.100866 (2015).

57 Debrabant, B. et al. Human longevity and variation in DNA damage response and repair: study of the contribution of sub-processes using competitive gene-set analysis. Eur J Hum Genet 22, 1131–1136, doi:10.1038/ejhg.2013.299 (2014).

58 Soerensen, M. et al. Human longevity and variation in GH/IGF-1/insulin signaling, DNA damage signaling and repair and pro/antioxidant pathway genes: cross sectional and longitudinal studies. Exp Gerontol 47, 379–387, doi:10.1016/j.exger.2012.02.010 (2012).

59 Chen, T. et al. A functional single nucleotide polymorphism in promoter of ATM is associated with longevity. Mechanisms of ageing and development 131, 636–640, doi:10.1016/j.mad.2010.08.009 (2010).

60 Piaceri, I., Bagnoli, S., Tedde, A., Sorbi, S. & Nacmias, B. Ataxia-telangiectasia mutated (ATM) genetic variant in Italian centenarians. Neurological sciences: official journal of the Italian Neurological Society and of the Italian Society of Clinical Neurophysiology 34, 573–575, doi:10.1007/s10072-012-1188-5 (2013).

61 Moskalev, A. A. et al. The role of DNA damage and repair in aging through the prism of Koch-like criteria. Ageing research reviews 12, 661–684, doi:10.1016/j.arr.2012.02.001 (2013).

62 Feng, Z., Hanson, R. W., Berger, N. A. & Trubitsyn, A. Reprogramming of energy metabolism as a driver of aging. Oncotarget 7, 15410–15420, doi:10.18632/oncotarget.7645 (2016).

63 White, R. R., & Vijg, J. Do DNA Double-Strand Breaks Drive Aging? Molecular cell 63, 729–738, doi:10.1016/j.molcel.2016.08.004 (2016).

64 Garinis, G. A., van der Horst, G. T., Vijg, J. & Hoeijmakers, J. H. DNA damage and ageing: new-age ideas for an age-old problem. Nature cell biology 10, 1241–1247, doi:10.1038/ncb1108-1241 (2008).

65 Fang, E. F. et al. NAD+ Replenishment Improves Lifespan and Healthspan in Ataxia Telangiectasia Models via Mitophagy and DNA Repair. Cell Metab 24, 566–581, doi:10.1016/j.cmet.2016.09.004 (2016).

66 Mills, K. F. et al. Long-Term Administration of Nicotinamide Mononucleotide Mitigates Age-Associated Physiological Decline in Mice. Cell Metab 24, 795–806, doi:10.1016/j.cmet.2016.09.013 (2016).

67 Zhang, H. et al. NAD(+) repletion improves mitochondrial and stem cell function and enhances life span in mice. Science 352, 1436–1443, doi:10.1126/science.aaf2693 (2016).

68 De Sandre-Giovannoli, A. et al. Lamin a truncation in Hutchinson-Gilford progeria. Science 300, 2055, doi:10.1126/science.1084125 1084125 [pii] (2003).

69 Eriksson, M. et al. Recurrent de novo point mutations in lamin A cause Hutchinson-Gilford progeria syndrome. Nature 423, 293–298, doi:10.1038/nature01629 nature01629 [pii] (2003).

70 Ghosh, S., Liu, B., Wang, Y., Hao, Q. & Zhou, Z. Lamin A Is an Endogenous SIRT6 Activator and Promotes SIRT6-Mediated DNA Repair. Cell reports 13, 1396–1406, doi:10.1016/j.celrep.2015.10.006 (2015).

71 Krishnan, V. et al. Histone H4 lysine 16 hypoacetylation is associated with defective DNA repair and premature senescence in Zmpste24-deficient mice. Proc Natl Acad Sci U S A 108, 12325–12330, doi:10.1073/pnas.1102789108 (2011).

72 Liu, B. et al. Depleting the methyltransferase Suv39h1 improves DNA repair and extends lifespan in a progeria mouse model. Nat Commun 4, 1868, doi:10.1038/ncomms2885 (2013).

73 Varela, I. et al. Accelerated ageing in mice deficient in Zmpste24 protease is linked to p53 signalling activation. Nature 437, 564–568, doi:nature04019 [pii] 10.1038/nature04019 (2005).

74 Endisha, H. et al. Restoring SIRT6 Expression in Hutchinson-Gilford Progeria Syndrome Cells Impedes Premature Senescence and Formation of Dysmorphic Nuclei. Pathobiology 82, 9–20, doi:10.1159/000368856 (2015).

75 The Selection and Use of Essential Medicines. World Health Organ Tech Rep Ser, vii–xv, 1–546 (2015).

76 Emami, J., Gerstein, H. C., Pasutto, F. M. & Jamali, F. Insulin-sparing effect of hydroxychloroquine in diabetic rats is concentration dependent. Can J Physiol Pharmacol 77, 118–123 (1999).

77 Razani, B., Feng, C. & Semenkovich, C. F. p53 is required for chloroquine-induced atheroprotection but not insulin sensitization. J Lipid Res 51, 1738–1746, doi:10.1194/jlr.M003681 (2010).

78 Schneider, J. G. et al. ATM-dependent suppression of stress signaling reduces vascular disease in metabolic syndrome. Cell Metab 4, 377–389, doi:10.1016/j.cmet.2006.10.002 (2006).

79 Yang, Y. P. et al. Application and interpretation of current autophagy inhibitors and activators. Acta Pharmacol Sin 34, 625–635, doi:10.1038/aps.2013.5 (2013).

80 Kimura, T., Takabatake, Y., Takahashi, A. & Isaka, Y. Chloroquine in cancer therapy: a double-edged sword of autophagy. Cancer Res 73, 3–7, doi:10.1158/0008-5472.CAN-12-2464 (2013).

81 Eisenberg, T. et al. Cardioprotection and lifespan extension by the natural polyamine spermidine. Nature medicine 22, 1428–1438, doi:10.1038/nm.4222 (2016).

82 Kenyon, C., Chang, J., Gensch, E., Rudner, A. & Tabtiang, R. A C. elegans mutant that lives twice as long as wild type. Nature 366, 461–464, doi:10.1038/366461a0 (1993).

83 Zhou, N., Xiao, H., Li, T. K., Nur, E. K. A. & Liu, L. F. DNA damage-mediated apoptosis induced by selenium compounds. The Journal of biological chemistry 278, 29532–29537, doi:10.1074/jbc.M301877200 (2003).

84 Zhang, Y. W. et al. Genotoxic stress targets human Chk1 for degradation by the ubiquitin-proteasome pathway. Molecular cell 19, 607–618, doi:10.1016/j.molcel.2005.07.019 (2005).

85 Ran, F. A. et al. Genome engineering using the CRISPR-Cas9 system. Nat Protoc 8, 2281–2308, doi:10.1038/nprot.2013.143 (2013).

86 Luo, B., Groenke, K., Takors, R., Wandrey, C. & Oldiges, M. Simultaneous determination of multiple intracellular metabolites in glycolysis, pentose phosphate pathway and tricarboxylic acid cycle by liquid chromatography-mass spectrometry. J Chromatogr A 1147, 153–164, doi:10.1016/j.chroma.2007.02.034 (2007).

